# Long-Range Enhancer Networks Gradually Emerge as Key Regulators During Human Cortical Neurogenesis

**DOI:** 10.1101/2024.07.24.605026

**Authors:** Lumeng Jia, Yingping Hou, Zejia Cui, Xin Luo, Jiali Duan, Jianbin Guo, Minglei Shi, Hebing Chen, Tingting Li, Fengyun Zhang, Bing Su, Lan Zhu, Cheng Li

## Abstract

The chromatin in the human brain cortex shows more enhancer-enhancer contacts than in macaques and mice, yet the organization of these contacts across cellular states and their mechanisms remain unclear. Here, we developed SCOPE-C, a cost-effective technique designed for mapping DNase I Hypersensitive Sites and their spatial interactions, with the potential for scalability to single-cell resolution. Employing SCOPE-C, we generated a comprehensive chromatin map detailing cis-regulatory elements in four cell types during neurogenesis in human, macaque, and mouse fetal cortex. This map introduces a model of long-range enhancer networks that regulate gene transcription during human cortical neurogenesis. In this model, CTCF-mediated loop extrusion forms ‘stripes’ of enhancer interaction hubs extending up to 10 Mb in human Excitatory Neurons. Enriched with human-specific enhancers and neuropsychiatric disorder-linked SNPs, these remarkably long-range networks predominantly control EN marker genes, suggesting their critical role in robust gene expression during human cortical neurogenesis.

## Background

The human brain has developed a larger and more complex structure compared to other species during the course of evolution, especially in the prefrontal cortex (PFC), which plays a crucial role in executing complex brain functions^1^. This expansion is primarily believed to be mediated by differences in spatiotemporal gene regulation rather than direct changes in protein-coding sequences^2^. Gene transcription regulation is considered a key mechanism in the development and evolution of the human brain^1,3–5^. Key elements in this regulatory landscape include cis-regulatory elements (CREs) such as enhancers and promoters, and trans-acting factors like transcription factors (TFs). For instance, recent studies have highlighted the role of retinoic acid in upregulating the expression of the *CBLN2* gene through its interaction with enhancer regions uniquely altered in the human brain^6,7^. Extensive omics studies have revealed that Human Accelerated Regions (HARs) and human gained enhancers shape the unique morphology of the human brain by promoting the expression of developmental genes^8–12^. Moreover, mutations in CREs in the human brain, especially those unique to humans, are closely related to neurological diseases and their progression^2,13–15^, making enhancers a hot topic of research in brain development^16–19^. However, linking enhancers to specific genes in the developing human brain is challenging. For instance, our previous work generated 3D and multi-omics data from the macaque fetal cortex. By integrating them with similar human datasets, we found that the human brain has more E-E interactions than those of macaques and mice^20^. Nevertheless, the high cell-type specificity of enhancers and their propensity for forming dynamic and complex chromatin spatial conformations with target genes make it exceptionally challenging to accurately delineate their regulatory roles and interactions with associated genes.

In terms of constructing cell-type specific 3D genomic maps, as technology has advanced, 3D genomics technologies have expanded to the single-cell level, even capable of simultaneously capturing 3D genome, methylation signals, or gene expression profiles in the same single cell^21–25^. However, these methods are usually not sufficient to map chromatin interactions between CREs at high resolution and are more suitable for analyzing larger chunks of chromatin interactions such as compartments or Topologically Associated Domains (TADs)^26–29^. To comprehensively capture interactions between CREs, these methods often require ultra-deep sequencing, which is not only costly but also demands high computational platforms^29^. Furthermore, existing methods for enriching CRE interactions usually require tens of thousands of starting cells to construct maps^30–32;^ Although the latest technologies attempt to reduce the required cell number to 1000^33^, the effect of formaldehyde crosslinking on the integration efficiency of Tn5 poses a challenge to comprehensively capturing the spatial interaction maps of CREs in low-input human brain samples.

With regard to interpret 3D genome data in the human brain, current studies mainly focus on short-distance interactions less than 1 mega base (Mb) within TADs^34–37^, while key regulatory events might occur over longer distances, even across multiple TADs, especially for developmentally important genes^38–40^. In addition, the mechanisms underlying the formation of transcription regulation-related loops remain unclear^41^, even though loop extrusion is widely recognized as a primary method for creating CTCF loops^42,43^, and the “chromatin stripes’’ are considered a manifestation of loop extrusion sliding^44^. Disrupting loop extrusion globally results in the extensive loss of CTCF loops^45^. However, Promoter-Enhancer (PE) interactions are generally stable, with only a minor impact on gene expression^45,46^. Recent research indicates that TFs-mediated condensates predominantly facilitate nested networks of PE interactions, especially for those active enhancers that are distantly located from their target genes^47^. Nevertheless, in HeLa cells, the acute depletion of cohesin drastically compromises long-distance (over 100 kb) promoter-anchored interactions^48^. Moreover, CTCF binding at promoters has been shown to enhance remote enhancer-dependent transcription across various cell types, underscoring the significance of CTCF and loop extrusion in transcriptional regulation.

To overcome these challenges, we developed SCOPE-C, a high-efficiency, low-cost sequencing technology that is not constrained by the initial number of cells and is capable of capturing spatial contacts between CREs more comprehensively. Using SCOPE-C, we constructed 212 DNA libraries from various cell lines (HeLa, K562, GM12878, NAMALWA, mESC, and mNSC) and four types of primary brain cells, representing different stages of neural genesis in the cerebral cortex of humans, macaques, and mice. We conducted a comprehensive analysis of the dynamic spatial interaction networks among CREs during cortical neurogenesis in fetal brains across various species, encompassing all interaction distance scales. Our findings revealed unique gene regulatory patterns in humans, elucidated their formation mechanisms, and highlighted their potential roles in human brain development.

## Results

### Develop SCOPE-C for detecting CREs and enriching their spatial contacts in capable of 1000 cells

SCOPE-C is a modified *in situ* Hi-C procedure^29^ designed for the efficient enrichment of open chromatin and their spatial interactions. Instead of utilizing sonication to fragment the chromatin after proximity ligation, SCOPE-C utilizes DNase I. This enzyme’s comparatively smaller size facilitates enhanced penetration and more precise cleavage within the nuclei of cross-linked cells, thereby ensuring robust information capture, even from low-input samples^49^. Additionally, for subsequent PCR amplification, we utilized gel extraction to collect library fragments, separating smaller fragments (160-200 bp) representing DHSs^49^ from larger fragments (200-500 bp) indicating spatial interactions among DHSs (**Figure 1A, Figure S1 and STAR Methods**). This step ensures that sufficient chromatin interaction data is gathered for analysis.

**Figure 1.**
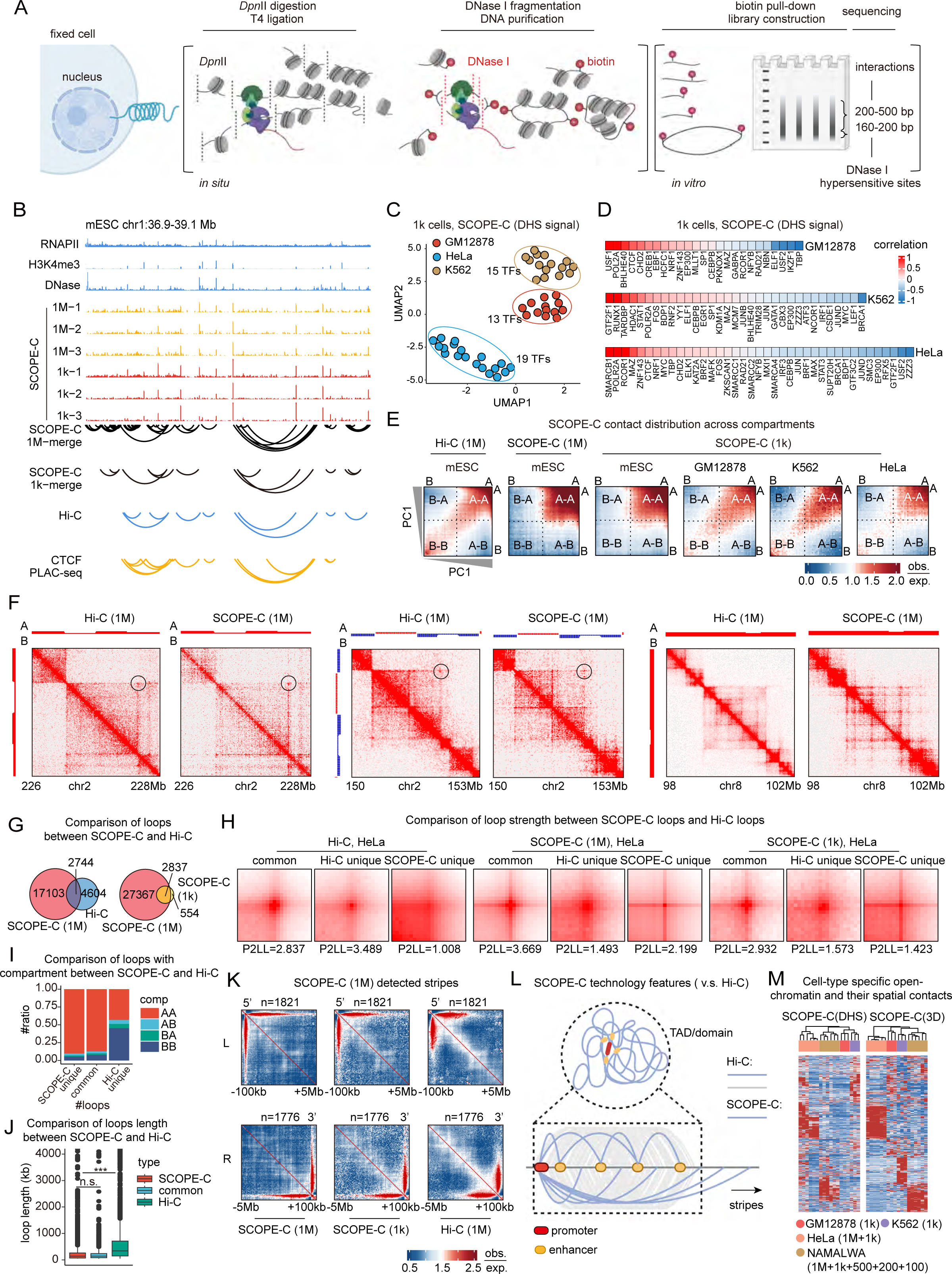
Develop SCOPE-C for detecting CREs and enriching their spatial contacts in at least 1 to 1000 cells. A. Schematic view of SCOPE-C showing the experimental procedure. B. Genome browser visualization of public ChIP-seq signals, along with SCOPE-C DNase I signals, chromatin loops identified from *in situ* Hi-C, CTCF PLAC-seq, and SCOPE-C in mESCs, using either 1 million or 1,000 cells. C. UMAP plot of all SCOPE-C libraries, profiled based on ChIP-seq signals localized within their DHS (DNase I Hypersensitive Site) peaks. Different colors are used to distinguish the cell types represented in each library. D. Heatmap displaying the generalized linear regression coefficients for DNA binding proteins within the SCOPE-C DHS peaks. E. Saddle plot indicating average contact enrichment between contacts of 100 kb bin arranged by their PC1 (shown on left and bottom): Lower left: B-B interactions; Upper right: A-A interactions; Upper left and lower right: A-B interactions. PC1 were called with Hi-C data of each cell types. F. Heatmap presentation of loops detected by Hi-C and SCOPE-C. G. Venn diagrams comparing loops identified by Hi-C and SCOPE-C across various input cell quantities. H. Aggregate Peak Analysis (APA) reveals the Hi-C and SCOPE-C reads enrichment surrounding SCOPE-C and Hi-C loops. I. Relationship between different types of SCOPE-C and Hi-C loops with AB compartments. J. Relationship between different types of SCOPE-C and Hi-C loops with loop length. K. Aggregate Stripe Analysis (ASA) displays SCOPE-C and Hi-C reads enrichment around SCOPE-C stripes in HeLa cells, each characterized by their distinct orientations. L. A cartoon model illustrating the preferential interactions captured by SCOPE-C compared to Hi-C. M. Heatmap presentation of both DHS signals and 3D SCOPE-C signals across various libraries, highlighting differences in cell types and the amount of input cells used, in human brain.

To assess DNase I’s ability to selectively target open chromatin, we sorted fresh and cross-linked cells from GM12878, K562, and HeLa cell lines, and conducted DNase-seq and SCOPE-C experiments at various DNase I concentrations (0.01, 0.03, 0.05, 0.07, and 0.1 U/uL). Analysis showed that 0.03 U/uL of DNase I effectively enriched DHSs near transcription start sites (TSSs). Peak analysis called from DNase-seq and small fragments of SCOPE-C, along with an assessment of chromatin states at these peak regions, also indicated that 0.03 U/uL DNase I preferentially cuts active regulatory elements. Thus, we confirmed that optimal digestion is achieved at 0.03 U/µL DNase I across various cell lines, regardless of cell type or fixation method (**Figure S2 and STAR Methods**).

To evaluate the effectiveness of SCOPE-C in enriching CREs and their interactions, as well as assessing the impact of using minimal cell inputs, we constructed libraries from 1 million and 1,000 cells in mESC, HeLa, K562, and GM12878 cell lines. Using HiC-Pro^50^ for alignment, we processed peaks for dangling ends and 160-200 bp valid pairs with MACS2^51^, and assessed interactions over 200 bp with FitHiC^52^ (**Figure 1B and STAR Methods**). We also integrated 25 GM12878, 33 K562, and 39 HeLa ChIP-seq datasets from ENCODE^53^, encompassing a variety of regulatory proteins. The comparative analysis showed that SCOPE-C peaks are clustered by cell type (**Figure 1C**) and are highly correlated with cell-type-specific TFs and structural proteins (**Figure 1D**). In comparisons within HeLa cells, SCOPE-C detected significantly more loops than Hi-C at identical sequencing depths and cell amount (19,847 *vs.* 7,348). Distinctively, loops unique to SCOPE-C and those shared with Hi-C displayed significant clustering within the A compartments, and also characterized by notably shorter interaction distances (**Figure 1E to 1J**) (*P <* 2.2e-16, Wilcoxon rank sum test), consistent with findings from previous studies^26,32^.

SCOPE-C was more effective at capturing long-range chromatin stripes^44^ (up to 5 Mb), due to its ability to eliminate background noise within TADs by distinguishing between hierarchical and functional interactions of CREs (**Figure 1K and 1L**). Finally, we expanded our experiments to include sorting fewer fixed cells (500, 200, and 100) in NAMALWA lines, and even tested SCOPE-C on as few as 12 single cells in mESC. By comparing the correlation between DHSs and interactions, we demonstrated SCOPE-C’s capacity as an effective 3D multi-omics technology for detailing cell-type specific chromatin spatial interactions, from 1000 cells to large populations and also its promising potential for extension to single-cell analyses (**Figure 1M, Figure S3, Figure S4 and STAR Methods**).

### Characterization of cell type-specific spatial contacts among CREs during human fetal cortical neurogenesis

To comprehensively capture CREs and their spatial contacts during human cortical development, we isolated four different types of cells (Radial Glias, RGs; Intermediate Progenitor Cells, IPCs; ENs and Interneurons, INs) representing different cell states during neurogenesis in the cortex from the PFC and visual cortex (V1) of the human fetal cortex during mid-gestation (19 gestational weeks), when neurogenesis and migration are at their peak^20^. Fluorescence-activated nuclei sorting (FANS) was used to sort these four types of cells from the fetal cortex. The gate, region, and cell proportions for these four types of cells were congruent with those of previous FANS experiments^36^, suggesting that the cells were isolated successfully **(Figure 2A and 2B, Figure S5A)**.

**Figure 2.**
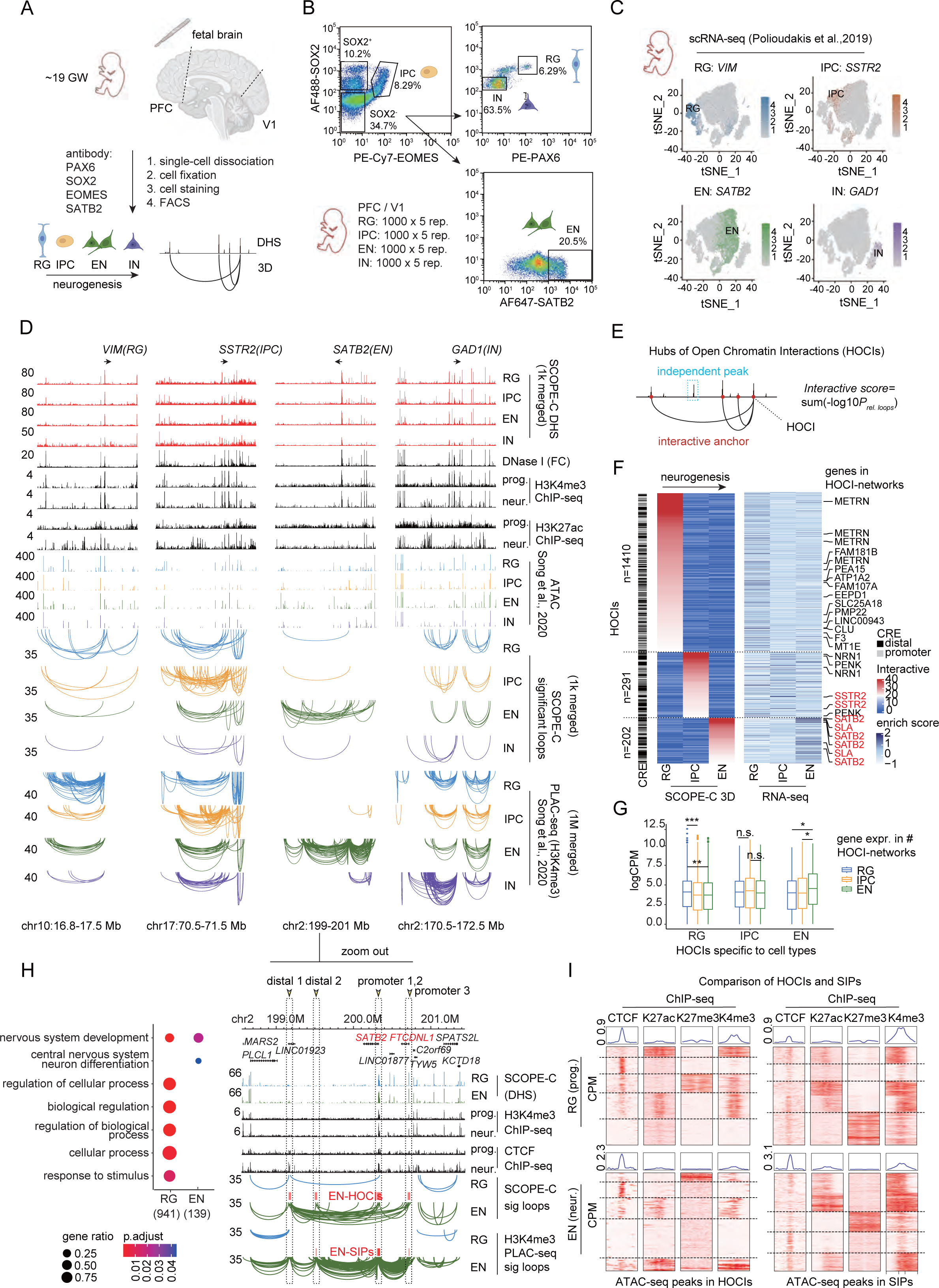
Characterization of neuron-specific spatial contacts among CREs during human fetal cortical neurogenesis. A. Schematic illustration of the sorting strategy detailing the antibodies employed to isolate specific human cortical neurons. B. FANS gating results of human cortex. C. Single cells from the human fetal cortex are depicted in t-SNE plot, with each cell color-coded according to the expression levels of the genes listed above. D. Graphical plots display the DHS (top and bottom) and 3D (middle and bottom) epigenomic landscapes, as characterized by SCOPE-C and supplemented with other public datasets (DNase-seq, ChIP-seq, ATAC-seq, PLAC-seq), surrounding marker genes for RG, IPC, EN and IN. A zoom in detailed view focused on *SATB2*, highlighting the HOCIs and SIPs. E. A cartoon model illustrating how to call HOCIs with SCOPE-C data. F. Heatmaps illustrate the types of cis-regulatory elements (CREs) on the left, the contact strengths in the middle, and HOCI-regulated gene expression on the right for SCOPE-C HOCIs, categorized by cell type specificity. The contact strengths are depicted by aggregating the log10-transformed false discovery rate (FDR) of loops, as derived from the FitHiC pipeline. In RG, IPC, and EN, we identified 1410, 291, and 202 cell-type specific HOCIs, respectively. G. Box plots showing the expression levels of genes regulated by HOCIs specific to RG, IPC and EN. H. Gene Ontology analysis showing the cell-type specific HOCI-regulated gene ontologies for RGs (941 genes) and ENs (139 genes). The color coding corresponds to enrichment scores calculated by gProfiler FDR, and the size indicates the ratio of genes. I. Heatmaps present the signal profiles for CTCF, H3K27ac, H3K27me3 and H3K4me3 around ATAC-seq peaks within HOCIs and super interactive promoters (SIPs) that are specific to RG, IPC and EN in the human fetal cortex.

To construct SCOPE-C libraries, we collected 3,000 to 10,000 cells from each type and used 1000 cells per type as input. We ensured 5 replicates for each cell type. In total, 62 SCOPE-C libraries were sequenced **(Table S1)**. As expected, SCOPE-C DHS signals were significantly enriched around TSSs and clustered by cell types, suggesting SCOPE-C reproducibly detected open chromatin signals in all types of cells across brain cortical regions **(Figure S6 and Figure S7)**. In order to compare open chromatin signals from PFC and V1, we combined SCOPE-C replicates and clustered the merged libraries. We observed that cells from distinct cortical regions tend to cluster together rather than cortical regions **(Figure S8)**. To comprehensively detect CREs and loops among them, we merged SCOPE-C data from PFC and V1 according to cell types.

In total, we identified 273,017, 326,670, 278,848, and 193,531 SCOPE-C peaks and 10,510, 8,292, 6,236, and 2,307 loops in RGs, IPCs, ENs, and INs. We downloaded the ATAC-seq data and H3K4me3 PLAC-seq data from the GEO database which were in line with our four cell types and sampling periods^36^ **(STAR Methods)**. The consistent peaks and loops between SCOPE-C, ATAC-seq and H3K4me3 PLAC-seq surrounding *VIM*, *SSTR2*, *SATB2* and *GAD1*, the well-known lineage-specific genes of RGs, IPCs, ENs, and INs, respectively^36^, suggesting the accuracy of cell type-specific DHSs and their spatial contacts in human fetal cortex **(Figure 2C and 2D)**. Furthermore, despite the consistent strength of their DHS signals across different cell types, these four genes establish various cell-type-specific loops **(Figure 2D)**. This suggests that chromatin spatial contacts among CREs are crucial for gene activation.

### Extensive Enhancers Interaction Hubs Regulate Lineage-Specific Gene Expression during Human Neurogenesis

Recent studies have demonstrated that gene expression is enhanced by active promoters with increased chromatin interactivity, particularly affecting lineage-specific genes^26,36^. To streamline our approach, we focused on CRE interaction networks rather than all loops, employing the concept of hubs of open chromatin interactions (HOCIs)^28^ and an interactive strength algorithm to define HOCIs as sets of open-chromatin regions (OCRs) with notably strong interactions **(Figure 2E and STAR Methods)**. These HOCIs form networks with their interacting regions. 762, 256, and 152 HOCIs were detected specific to human RGs, IPCs, and ENs, respectively. Notably, we did not calculate HOCIs in INs as these cells do not directly differentiate from cortical progenitors. Unlike the other three cell types, INs were not sequenced as deeply to explore the spatial dynamics of CRE interactions during the differentiation process from neural progenitors to radial neurons.

To further characterize these HOCIs, we analyzed genomic regions and ChIP-seq signals of developmental neurons from the ENCODE^53^ database around these HOCIs. To prevent bias in detecting DREs with SCOPE-C, we conducted our analysis based on ATAC-seq peaks within HOCIs. We found that these HOCIs are identifiable as active promoters, enhancers, and insulators, distinct from the previously reported super-interacting promoters (SIPs)^36^ primarily marked by the H3K4me3 ChIP signal **(Figure 2D and 2I)**. Of all HOCIs, 39% aligned with gene promoters (promoter HOCIs), while 61% associated with distal regions (distal HOCIs) **(Figure 2F)**.

By summing up the log-transformed FDR of SCOPE-C peaks and ATAC-seq peaks within each HOCI-regulated gene promoter, respectively, we calculated chromatin openness for each promoter of HOCI-regulated genes. We found that either SCOPE-C or ATAC-seq supported that chromatin openness at these HOCI-regulated gene promoters remained stable across RGs, IPCs and ENs **(Figure S9)**, confirming that most emerging OCRs during development are found in distal regions rather than gene promoters^54,55^.

To link the HOCIs with gene expression, we considered genes regulated by HOCIs if their promoters overlapped with or were proximal to HOCIs, or if their promoters interacted with distal HOCIs, as indicated by SCOPE-C loops. RNA-seq data analysis showed a positive correlation between HOCI interaction strength and regulated gene expression levels in RGs and ENs (**Figure 2F and 2G**) (stripes in RGs: *P_(RG v.s. IPC)_* =3.644e-05, *P_(RG v.s. EN)_ =* 0.000181; stripes in IPCs: *P_(IPC v.s. RG)_* = 0.1202, *P_(IPC v.s. EN)_ =* 0.1817; stripes in ENs: *P_(EN v.s. RG)_* = 0.009006, *P_(EN v.s. IPC)_ =* 0.01005; Wilcoxon rank sum test). Gene Ontology (GO) analysis confirmed that HOCI-regulated genes are lineage-specific (**Figure 2H**). We conclude that in addition to confirming that lineage genes with promoters involved in extensive interactions with regulatory elements exhibit high expression levels, as previously concluded, we have also revealed that lineage genes interacting within the hubs of extensive enhancers similarly demonstrate elevated expression levels.

### Dynamic CRE Interaction Networks Involved in CTCF-Mediated Long-Range Loop Extrusion for Gene Transcription Regulation

We aimed to understand why the human brain exhibits more E-E interactions. As loops can pre-exist before gene expression and remain stable throughout development^21,56,57^, distinguishing whether spatial interactions or other factors regulate gene transcription presents a challenge. Consequently, we focused on dynamic HOCIs (dHOCIs) and their regulatory networks, which exhibit significant changes during development.

Previously, we found that HOCI regulation could distinctly differentiate gene expression levels between radial glia (RGs) and excitatory neurons (ENs). To investigate the association between chromatin loops and gene expression, we identified dynamic loops and characterized dHOCIs as chromatin regions showing the greatest variability in interactivity during the RG-to-EN transition. We categorized these dHOCIs into two types: EN-gained and EN-lost, and linked dHOCI-regulated genes to these categories (**Figure 3A and 3B, STAR Methods**). We observed that expression levels of dHOCI-regulated genes were positively correlated with their interaction strength (**Figure 3C and 3D**) (*P_EN-gained_*=0.0002072 and *P_EN-lost_=* 3.293e-05, Wilcoxon rank sum test). Notably, EN-gained dHOCIs (102) outnumbered EN-lost dHOCIs (60), despite more HOCIs being present in RGs than in ENs (**Figure 2F and Figure 3B**). Moreover, dynamic interaction scores in EN-gained dHOCIs were significantly higher than those in EN-lost (**Figure 3B**), indicating that newly acquired CRE interaction networks in ENs play a critical role in regulating gene expression.

**Figure 3.**
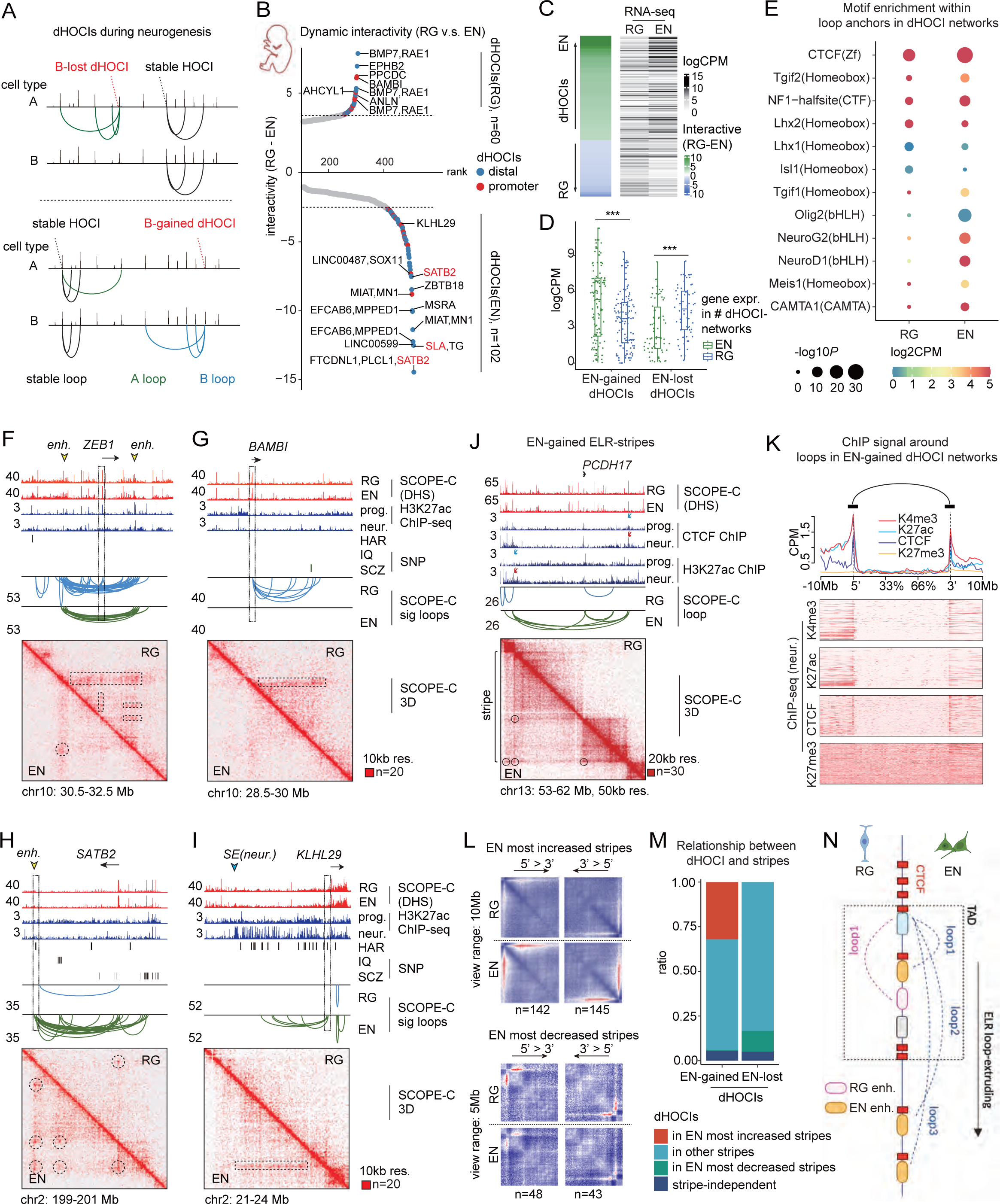
Loop Extrusion Drives CRE Interaction Networks for Gene Transcription Regulation. A. A cartoon model illustrating how to call dHOCIs with SCOPE-C data during cell development. B. Dynamic HOCIs (dHOCIs) have been prioritized based on their differential interaction scores of HOCI-related loops in RGs and ENs. dHOCIs are highlighted in red for proximal CERs and blue for distal CREs. A total of 60 SIOs for RG and 102 dHOCIs for EN were ultimately identified. C. Heatmaps showing the contact strengths on the left, and dHOCI-regulated gene expression on the right for RG to EN. D. Box plots showing the expression levels of genes regulated by dHOCIs specific to RG to EN. E. Dot plot illustrates the enrichment of motifs within SCOPE-C peaks at anchors of dHOCI-related loops. The colors indicate the gene expression levels of the respective TFs, while the sizes correspond to the P-values obtained from HOMER analysis. F-J. The plot visualizes dHOCIs from RG to EN. The H3K27ac ChIP-seq signal highlights active enhancers in both neuronal progenitors and differentiated neurons. The plot also incorporates Human Accelerated Regions (HARs) and Single Nucleotide Polymorphisms (SNPs) linked to neuropsychiatric disorder. K. Average signal intensity profiles for H3K4me3, H3K27ac, CTCF, and H3K27me3 ChIP-seq reads distributed around the anchors of loops associated with EN-gained dHOCIs. L. Aggregate Stripe Analysis (ASA) display SCOPE-C reads enrichment around dynamic stripes from human RGs to ENs, each characterized by their distinct orientations. M. Relationship between dHOCIs and stripes. N. A cartoon model illustrating long-range loop extruding based CRE interaction formation.

To enrich our understanding of CRE interactions, particularly for lineage-specific genes, we explored potential mechanisms mediating dHOCI networks. First, we analyzed the motif enrichment of SCOPE-C peak within dHOCI and associated loop anchors. We found that CTCF was consistently and strongly enriched in both EN-gained and EN-lost dHOCI networks, supported by ChIP-seq data from ENCODE (**Figure 3E**). Furthermore, we discovered that H3K4me3 and H3K27ac ChIP signals were highly enriched in loop anchors, while H3K27me3 signals were absent (**Figure 3K**), further supporting the involvement of active CREs in dHOCI networks. Since CTCF-associated loops are often mediated by loop extrusion, we used SCOPE-C interactions to create heatmaps near dHOCI-regulated genes, revealing that these loops indeed coincide with stripes, often considered a result of loop extruding (**Figure 3F to 3J**). This suggests that interactions between genes and active CREs in the neural lineage of the brain are mediated by CTCF-mediated loop extruding.

Interestingly, we found that HOCI-related loops and stripes newly acquired in ENs were significantly longer than those specific to RGs, with stripes in ENs reaching up to 10 Mb and spanning across TAD boundaries (**Figure 3J, 3L and 3N**). Moreover, compared to dHOCIs lost in ENs, those gained in ENs were more likely to co-occur with dynamically emerging stripes in ENs, despite a considerable number of EN-gained dHOCIs being associated with stable stripes from RG to EN (**Figure 3M**). Recent research suggests that TAD boundary stacks drive long-distance interactions between some developmental important genes and their enhancers across TADs^40^. However, we found that in addition to TAD stacks, there are CREs independent of TAD boundaries that organize their loops through loop extruding to regulate gene expression (**Figure 3I**).

### Human-Specific Enhancers Are Enriched in Dynamic CRE Interaction Networks of Humans

To examine the distribution and function of dHOCI-associated CRE networks across species, we utilized SCOPE-C analysis on fetal cortical tissues from macaques (*Macaca mulatta*) and mice (*Mus musculus*) during equivalent gestational periods (macaque: 80 days; mouse: 14.5 days)^20,58^. We sorted 3,000 to 6,000 cells from RGs, IPCs, and ENs using FACS strategy in both species (**Figure 4A, Figure S5B and S5C**). For each cell type, 1,000 cells were used to construct SCOPE-C libraries, achieving 3 to 5 replicates (**Table S1**). We assessed the quality of the data and identified chromatin loops and peaks specific to each cell type, similarly to our analysis in the human brain (**Figure 4B, Figure S10 and Figure S11**).

**Figure 4.**
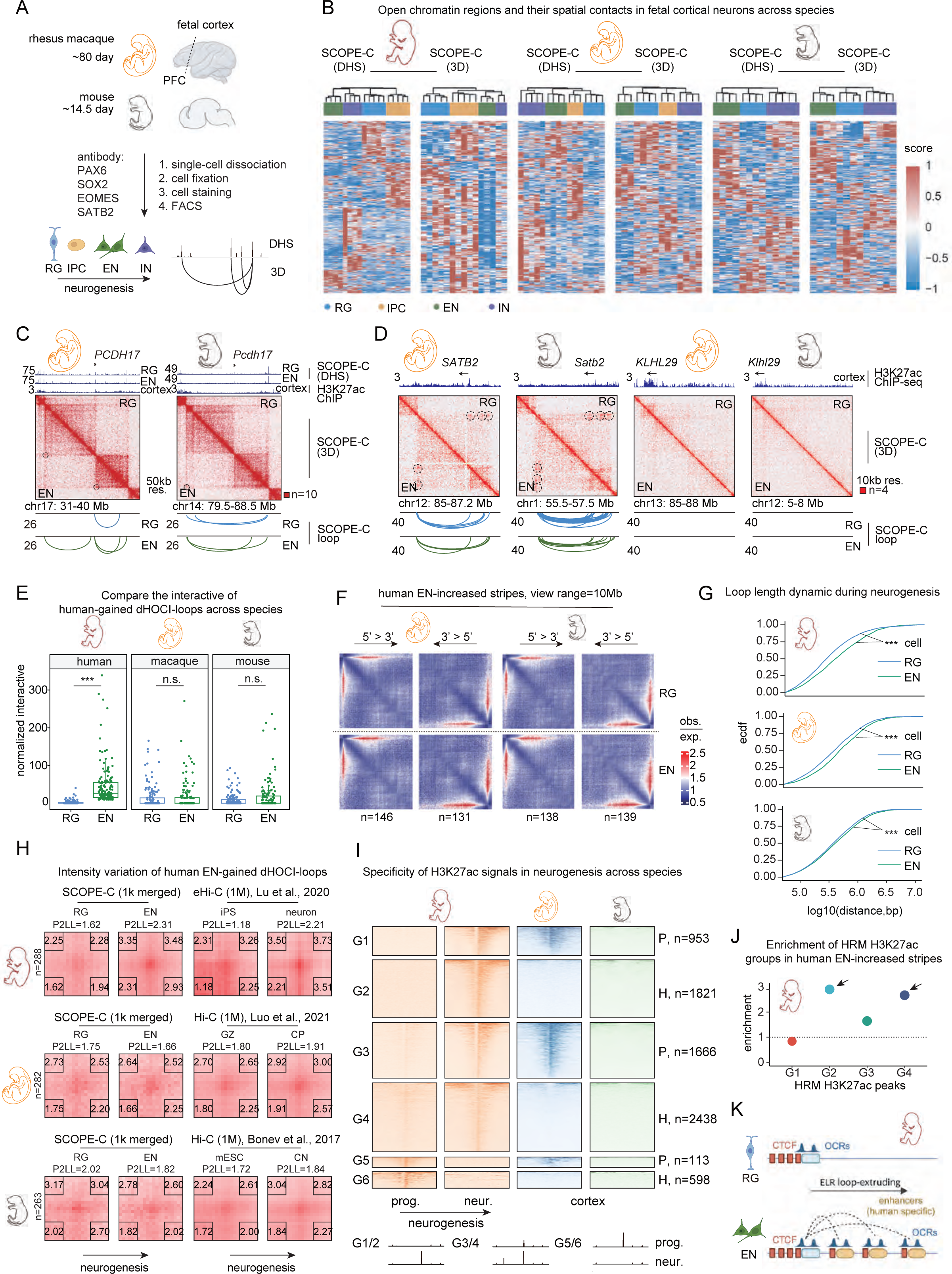
Dynamic stripes in CRE interaction networks unique to humans organize interactions with human-specific enhancers. A. Schematic illustration of the sorting strategy detailing the antibodies employed to isolate specific neuron types in macaque and mouse fetal brain. B. Heatmap presentation of both DHS and 3D SCOPE-C signals across various libraries, highlighting differences in neurons, spanning macaques and mice. C-D. Plots illustrate the interactions around the *SATB2*, *KLHL29* and *PCDH17* genes in macaque and mouse, corresponding to Figure 3g, j-k. Curves indicate significant loops detected by SCOPE-C, with colors distinguishing the cell types: royal blue for RG and forest green for EN. E. Comparing the interactive strength of human-EN-gained dHOCI-loops across species. F. Aggregate Stripe Analysis (ASA) showing enrichment of SCOPE-C reads from macaques and mice around human EN-gained dHOCIs in RGs and ENs across various species, each with distinct orientations. G. Comparation of loop length between SCOPE-C loops in RGs and ENs across species. H. Aggregate Peak Analysis (APA) reveals the reads enrichment surrounding human EN-gained dHOCI-related loops in RGs (or progenitors) and ENs (or neurons) across species, corroborated by SCOPE-C and public Hi-C datasets. I. H3K27ac peaks across species are analyzed, with grouping based on their emergence in specific species and cell types. ‘H’ denotes peaks exclusive to humans, while ‘P’ indicates those found only in primates. J. Scatter plot showing the enrichment of H3K27ac groups for anchors of human EN-gained dHOCI-related loops. K. A cartoon model illustrating the formation of long-range loop extrusion-based CRE interactions that organize human-specific enhancers.

Additionally, by examining open chromatin pairs and loops around those dHOCIs shown in Figure 3H, 3I, and 3J, we found that these dHOCIs are human specific **(Figure 4C and 4D)**. Based on our SCOPE-C data, human ENs exhibit a greater loop strength with loops linked to EN-gained-dHOCIs compared to human RGs. Moreover, homologous loops of these dHOCI-associated loops are specific to human cortical neurogenesis **(Figure 4E and 4H)** (Figure 4E, human: *P_(EN v.s. RG)_* < 2.2e-16, macaque: *P_(EN v.s. RG)_ =* 1; mouse: *P_(EN v.s. RG)_ =* 0.82; Wilcoxon rank sum test). This conclusion can also be corroborated by publicly available Hi-C data **(Figure 4H)**. In contrast, chromatin stripes in macaques and mice remained consistent across cell types without notable dynamic alterations (**Figure 4F**). Despite a general increase in loop length in ENs across all species, the magnitude of change was more pronounced in humans (**Figure 4G**).

To further explore the functional impact of chromatin stripes and dHOCI-related loops in human ENs, we analyzed H3K27ac ChIP-seq data from the GEO database (**STAR Methods**), documenting changes during neuron development. We identified six groups of H3K27ac peaks (G1 to G6) based on their activity during differentiation and presence across species (**Figure 4I**). Specifically, using homologous loops from human, macaque, and mouse (**STAR Methods**), we analyzed the enrichment of H3K27ac peaks (from G1 to G4) at these loop anchors. We observed a significant enrichment of human EN-specific H3K27ac peaks (G2 and G4) at these homologous loop anchors (**Figure 4J**), indicating that these human EN-acquired dHOCIs are associated with human-specific enhancers (**Figure 4K**).

It is well-established that enhancers unique to the human brain are densely populated with HARs and single nucleotide polymorphisms (SNPs) linked to neuropsychiatric disorders^58–60^. To further explore the potential relationship between these human-specific enhancers, particularly HARs, and the dHOCIs acquired by human ENs. We identified a series of HARs enriched within the EN-gained stripes involving the *KLHL29* gene (**Figure 3I**). ChIP-seq data supported the co-localization of these HARs with active enhancers in neuronal cells (**Figure 3I**). The KLHL29 gene is implicated in Bardet-Biedl Syndrome 7, marked by intellectual disabilities or developmental delays, highlighting its critical role in human brain evolution.

### Potential Role of Loop-Extrusion-Driven Long-Range Enhancer Interactions in Robust Gene Activation

Previous studies at the cellular population level of the human fetal cortex have demonstrated that human-specific loops primarily engage in E-E interactions, which enhance the expression levels of their regulated genes^15^. We discovered that enhancers regulated by these human-specific PE interactions are predominantly unique to humans and particularly relevant to cells (**Figure 4H to 4K**). We therefore further investigated whether the expression levels of genes regulated by these human EN-specific PE interactions differ across species. Public RNA-seq datasets from developing human and mouse fetal cortex were utilized for this purpose (**STAR Methods** and **Figure S12**). A differential gene expression analysis was performed to contrast brain responses across species. This analysis revealed that genes regulated by human EN-gained dHOCIs tend to show more pronounced expression in human ENs (**Figure 5A and 5B**). Although overall gene expression is upregulated in human ENs compared to mouse ENs, not all genes exhibit higher expression in humans relative to mice (**Figure 5C and 5D**) (Figure 5C, *P*=0.028, Wilcoxon rank sum test). For instance, *SATB2* and *KLHL29*, two representative genes regulated by human EN-gained dHOCIs, are shown in Figure 3J and 3K. Recent insights suggest that ultra-long-range enhancer networks, potentially based on phase separation, have a multi-layered nested structure that provides robust gene expression necessary to overcome genetic variations such as SNPs^61^. This supports the hypothesis that these chromatin stripe-mediated dHOCI networks not only upregulate gene expression but also contribute to robust gene activation. The strong enrichment of SNPs within loops associated with EN-gained dHOCIs reinforces this idea (**Figure 5E**). Further analyses of genes regulated by human EN-gained dHOCIs revealed a significant enrichment of transcription factors (TFs) and DNA-binding proteins (**Figure 5F**; Chi-square test, P=1.57e-7). Protein-protein interaction (PPI) analyses suggest potential synergistic interactions among these TFs, which, along with other TFs related to neurodevelopment, collaboratively regulate gene expression crucial for brain development **(Figure 5G)**.

**Figure 5.**
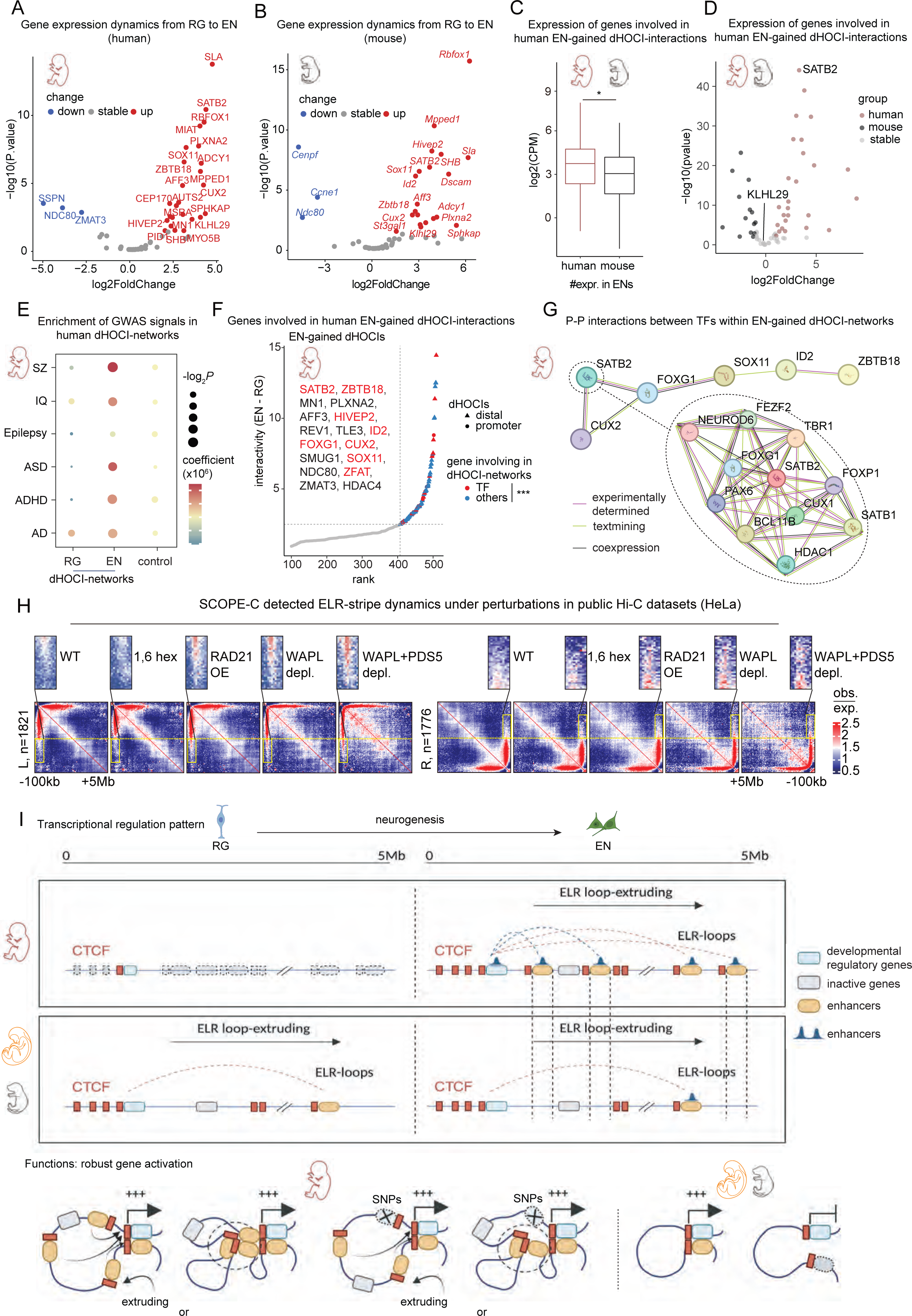
dHOCI-associated stripes contribute to more than gene activation. A-D. Volcano plots illustrate the distinct gene expression levels between ENs and mouse ENs across species. E. Dot plots depict the enrichment of SNPs in SCOPE-C peaks within dHOCI-related loops from RG to ENs. The color scale indicates the coefficients, and the size of each dot reflects the log10-transformed P values for dHOCI enrichment. F. Human EN-gained dHOCIs have been prioritized based on their differential interaction scores of HOCI-related loops in RGs and ENs. dHOCIs are highlighted in red for their regulated genes are TFs. G. PPI networks illustrate the interactions among transcription factors regulated by EN-gained dHOCIs. H. Aggregate Stripe Analysis (ASA) display Hi-C reads enrichment around SCOPE-C stripes in HeLa cells. I. Cartoon models hypothesize the function and mechanism of loop extrusion-driven human-specific dHOCIs for robust gene activation.

To further demonstrate that these chromatin stripe-mediated dHOCI networks are indeed driven by loop extrusion, we collected various Hi-C data from HeLa cells (**STAR Methods**), both wild-type and those with disrupted or enhanced phase separation and loop extrusion. By comparing the supported intensities of 314 most observed SCOPE-C stripes (5’ > 3’: 189; 3’ > 5’: 125, see method) in HeLa under different conditions, we found that extending the sliding distance of loop extrusion (WAPL depletion, WAPL+PDS5 depletion) or increasing cohesin loading on the chromatin (RAD21 overexpression) elongated the stripes. Conversely, disruptions to phase separation by 1,6 Hex did not significantly change the length of stripes. This indicates that these long-range stripes are indeed driven by loop extrusion (**Figure 5H**).

## Discussion

In this study, we develop SCOPE-C, a cost-effect 3D multi-omics tool designed to simultaneously enrich open chromatins and their spatial contacts using as few as 1,000 input cells. Such advancements address the challenge of limited cell availability in clinical samples. With this map, we introduce a loop extrusion-based model aimed at regulating robust gene activation during human cortical neurogenesis (**Figure 5I**). By analyzing dynamic CRE interaction networks, we extended the knowledge of cell-type-specific promoter-associated spatial contacts during human fetal cortical development^36^. This not only fills gaps in E-E interactions but also focuses on human-specific dynamic interaction networks and their regulatory genes across species. Moreover, previous studies have reported that redundant enhancers robustly activate target genes through a phase separation-mediated nested model^47^. Our model extends this mechanism. We propose that during human cortical neurogenesis, loop extrusion facilitates spatial interactions among PEs. Additionally, while past research often suggested that PE interactions are short-range, typically confined within TADs, our new techniques and analytical framework facilitate the discovery of extremely long-range interactions between CREs, spanning regions up to 10 Mb in ENs. This indicates that chromatin stripes, previously reported to exceed 1Mb in length and spanning TADs, and associated with super enhancers, indeed play a role in gene transcription regulation. In our model, we propose that extruding-mediated PE interactions may be transient. During the extruding process, gene promoters sequentially interact with redundant enhancers (**Figure 5I**). This finding is supported by recent live-cell imaging studies, which demonstrate that super-enhancers interact with Sox2. During these interactions, condensates, promoters, and enhancers transiently come together under the influence of loop extrusion to control gene bursting. Besides, human-specific enhancers and SNPs are enriched in these dynamic CREs interaction networks. This interaction pattern helps overcome genetic variations, ensuring consistent gene activation.

Furthermore, our model deepens the understanding of how CTCF regulates critical developmental genes. Our results suggest that the concurrent presence of loops and stripes within CREs is a potential indicator of high gene expression. This could be because stripes represent rapid, continuous sliding of loops, increasing the frequency of CRE contacts. Recent live-cell imaging technologies have reported that isolated Hi-C loops are observed only infrequently and in a minority of cells^59^, suggesting that Hi-C loops alone may not support the universality and persistence of spatial contacts in cell populations. Previous reports have indicated that after globally removing and then reconstructing loops, actively regulated elements, especially those involving super-enhancers, tend to rebuild more quickly, with visible stripes^45^. We believe that the co-occurrence of stripes and loops in spatial interactions may be more common and persistent across cell populations. This hypothesis may bridge the gap between sequencing-based methods and live-cell imaging in identifying loop structures, offering a new perspective on how chromatin interactions regulate gene transcription.

However, our study has limitations. Single-cell RNA sequencing studies have uncovered new cell types and subtypes in the human cortex^58,60,61^. FACS sorting for nuclei isolation may prove inadequate for fine cellular distinction. It did not counteract cortical cell sub-clusters and cell ratio influences across species. Moreover, the stripe identified in the mini-bulk cell populations does not definitively indicate its presence at the single-cell level, as it could be an artifact of averaging across the cell population. We believe that in the future, by integrating combinatorial labeling methods, SCOPE-C will be extended to single-cell analysis. With the use of high-throughput single-cell sequencing technologies, we will attempt to further address this lingering question.

In summary, our research sheds light on the intricate chromatin spatial contacts of the human genome, underscoring the critical role of dynamic chromatin contacts and their long-range extrusion processes in gene regulation. The correlation of these genomic characteristics with human cortical neurogenesis and neurological disorders provides critical insights into brain function and enhances our understanding and potential diagnosis of these conditions.

## Data and Code availability

All datasets used in this study are available at the Genome Sequence Archive for Human (GSA number: HRA007714) under controlled access and GEO (accession number: GSE267089).

## Acknowledgements

This work was supported by National Key Research and Development Program of China (2021YFA1100300) and National Natural Science Foundation of China (32288102, 32025006). Part of the data analysis was performed on the High-Performance Computing Platform of the Center for Life Sciences, Peking University. We thank the flow cytometry Core at National Center for Protein Sciences at Peking University, particularly Jia Luo and Liying Du, for technical help.

## Author contributions

C.L., L.M.J., conceived the study. C.L., L.Z., B.S. supervised the study. L.M.J. and Y.P.H. performed experiments. L.M.J. and Z.J.C. performed computational analysis and X.L., J.B.G., J.L.D., F.Y.Z. collected the fetal cortex samples from human, macaque and mouse, respectively. L.M.J. wrote the manuscript. M.L.S., H.B.C., T.T.L., offer the suggestions for the study.

## Extended Data Figure Legend

**Figure S1. Gel analysis of determining the optimal concentration of DNase I in SCOPE-C library construction and gel cutting strategy for sequencing**

The concertation titration of DNase I was initiated ranging from a to i, representing 0, 0.003, 0.005, 0.007, 0.01, 0.03, 0.05, 0.07, 0.1 U/μL, respectively

A. The gel analysis of SCOPE-C library at DNase I concentrations ranging from a to i in GM12878.

B. The gel analysis of SCOPE-C library at DNase I concentrations ranging from a to i in K562.

C. The gel analysis of SCOPE-C library at DNase I concentrations ranging from a to i in HeLa.

**Figure S2. Determining the optimal concentration of DNase I in SCOPE-C experiments**

A. The distances from the peaks to the TSS (transcription start sites) at different DNase I concentrations in DNase-seq experiments with different human cell lines (GM12878, K562, HeLa).

B. The chromHMM annotation of peaks at different DNase I concentrations in DNase-seq experiments with 3 types of human cells.

C. The distances from the peaks to the TSS (transcription start sites) at different DNase I concentrations in SCOPE-C experiments with different human cell lines (GM12878, K562, HeLa).

D. The chromHMM annotation of peaks at different DNase I concentrations in SCOPE-C experiments with 3 types of human cells.

**Figure S3. SCOPE-C accurately identifies DHS open-chromatin peaks in bulk/mini-bulk/single cells**

A. Genome browser showing public ChIP-seq peaks and SCOPE-C DHS peaks in NAMALWA with 1000, 500, 200 and 100 cell number.

B. Genomic features results of SCOPE-C DHS peaks under different cell volumes.

C. TSS plot showing reads distribution of SCOPE-C with differing starting cell numbers and public DNase-seq around TSS.

D. Pearson correlation between SCOPE-C DHS peak signal under different cell volumes.

E. Venn diagram showing the highly coincident SCOPE-C peaks called in NAMALWA with differing starting cell number.

F. Genome browser showing single-cell SCOPE-C data in mESC.

G. TSS plot in different single cell repeats.

H. Pearson correlation between million cells, 1000 cells, single-cell SCOPE-C DHS signal and public DNase-seq peak signal in mESC. Control is the SCOPE-C data from mouse neural stem cell (NSC) lines (*i.e.* ctrl-NSC)

**Figure S4. SCOPE-C faithfully capture the key feature of genome organization in bulk/mini-bulk/single cells**

A. mESC chromatin contact matrices of SCOPE-C (under the diagonal) and *in situ* Hi-C (above the diagonal) at successive zoom-in views.

B. Scatterplots comparing the eigenvector computed from SCOPE-C versus *in situ* Hi-C of mESC. PCC, Pearson correlation coefficient.

C. Scatterplots comparing the directionality index computed from SCOPE-C versus *in situ* Hi-C of mESC. PCC, Pearson correlation coefficient.

D. AB compartment comparison of chromosome 1 computed from SCOPE-C under different cell volumes in mESC.

E. TAD structure comparison of chromosome 2 (chr2:52-58 Mb) computed from SCOPE-C under different cell volumes in mESC.

**Figure S5. Representative contour plots depicting FACS gating strategy**

A. FANS gating strategy of human primary visual (V1) cortex.

B. FANS gating strategy of macaque cortex.

C. FANS gating strategy of mouse cortex.

**Figure S6. Quality control results for human fetal brain (PFC) SCOPE-C data**

A. SCOPE-C DHS signals of RG cells enrichments at TSS.

B. SCOPE-C DHS signals of IPC cells enrichments at TSS.

C. SCOPE-C DHS signals of EN cells enrichments at TSS.

D. SCOPE-C DHS signals of IN cells enrichments at TSS.

E. Clustering results of Pearson correlation between SCOPE-C DHS signals for the four cell types.

F. PCA dimensionality reduction results for SCOPE-C 3D signals for the four cell types.

**Figure S7. Quality control results for human fetal brain (V1) SCOPE-C data**

A. SCOPE-C DHS signals of RG cells enrichments at TSS.

B. SCOPE-C DHS signals of IPC cells enrichments at TSS.

C. SCOPE-C DHS signals of EN cells enrichments at TSS.

D. SCOPE-C DHS signals of IN cells enrichments at TSS.

E. Clustering results of Pearson correlation between SCOPE-C DHS signals for the four cell types.

F. PCA dimensionality reduction results for SCOPE-C 3D signals for the four cell types.

**Figure S8. Comparison of human prefrontal cortex (PFC) and primary visual (V1) DHS signals**

A. Feature distribution results of DHS peaks in different cell types of PFC and V1.

B. The intersection between DHS peaks from different cell types of PFC and V1.

C. Clustering results of Pearson correlation between DHS signals in different cell types of PFC and V1.

D. PCA dimensionality reduction results for DHS signals in different cell types of PFC and V1.

**Figure S9. The chromatin openness at HOCI-regulated gene promoters in SCOPE-C and ATAC-seq data**

**Figure S10. Quality control results for macaque fetal brain SCOPE-C data**

A. SCOPE-C DHS signals of RG cells enrichments at TSS.

B. SCOPE-C DHS signals of IPC cells enrichments at TSS.

C. SCOPE-C DHS signals of EN cells enrichments at TSS.

D. SCOPE-C DHS signals of IN cells enrichments at TSS.

E. Clustering results of Pearson correlation between SCOPE-C DHS signals for the four cell types.

F. PCA dimensionality reduction results for SCOPE-C 3D signals for the four cell types.

**Figure S11. Quality control results for mouse fetal brain SCOPE-C data**

A. SCOPE-C DHS signals of RG cells enrichments at TSS.

B. SCOPE-C DHS signals of EN cells enrichments at TSS.

C. SCOPE-C DHS signals of IN cells enrichments at TSS.

D. Clustering results of Pearson correlation between SCOPE-C DHS signals for the three cell types.

E. PCA dimensionality reduction results for SCOPE-C 3D signals for the three cell types.

**Figure S12. Principal component analysis results of “meta-cell” datasets composed of two groups of single-cell transcriptome data derived from human and mouse**

**Table S1:** Data samples and library information. In addition to the single-cell libraries, there are two other types of libraries: short-fragment (160-200 bp) libraries and long-fragment (200-500 bp) libraries, totaling 92 libraries.

|  | Cell lines | Bulk | 1000 | 500 | 200 | 100 | single cell |
| --- | --- | --- | --- | --- | --- | --- | --- |
| Human | HeLa | 3 | 5 | / | / | / | / |
|  | GM12878 | / | 5 | / | / | / | / |
|  | K562 | / | 5 | / | / | / | / |
|  | NAMALWA | / | 5 | 2 | 2 | 2 | / |
| Mouse | ESC | 3 | 3 | / | / | / | 10 |
|  | NSC | 3 | 3 | / | / | / | / |

**Table S2:** Data samples and library information. Different primary cell types were sorted from human, macaque, and mouse fetal brain samples, and all were subjected to SCOPE-C experiments using 1000 cells, with each sample containing two libraries: short-fragment (160-200 bp) libraries and long-fragment (200-500 bp) libraries, totaling 128 libraries.

|  | Primary samples | Human-PFC | Human-V1 | Macaque | Mouse |
| --- | --- | --- | --- | --- | --- |
| 1000 cells | RG | 5 | 4 | 4 | 4 |
|  | IPC | 5 | 4 | 4 | 1 |
|  | EN | 4 | 4 | 4 | 4 |
|  | IN | 4 | 4 | 4 | 5 |

**Table S3:**
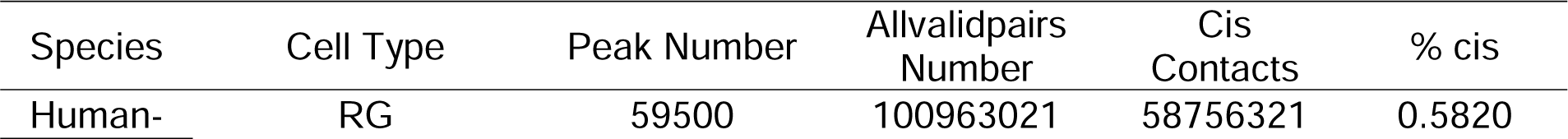
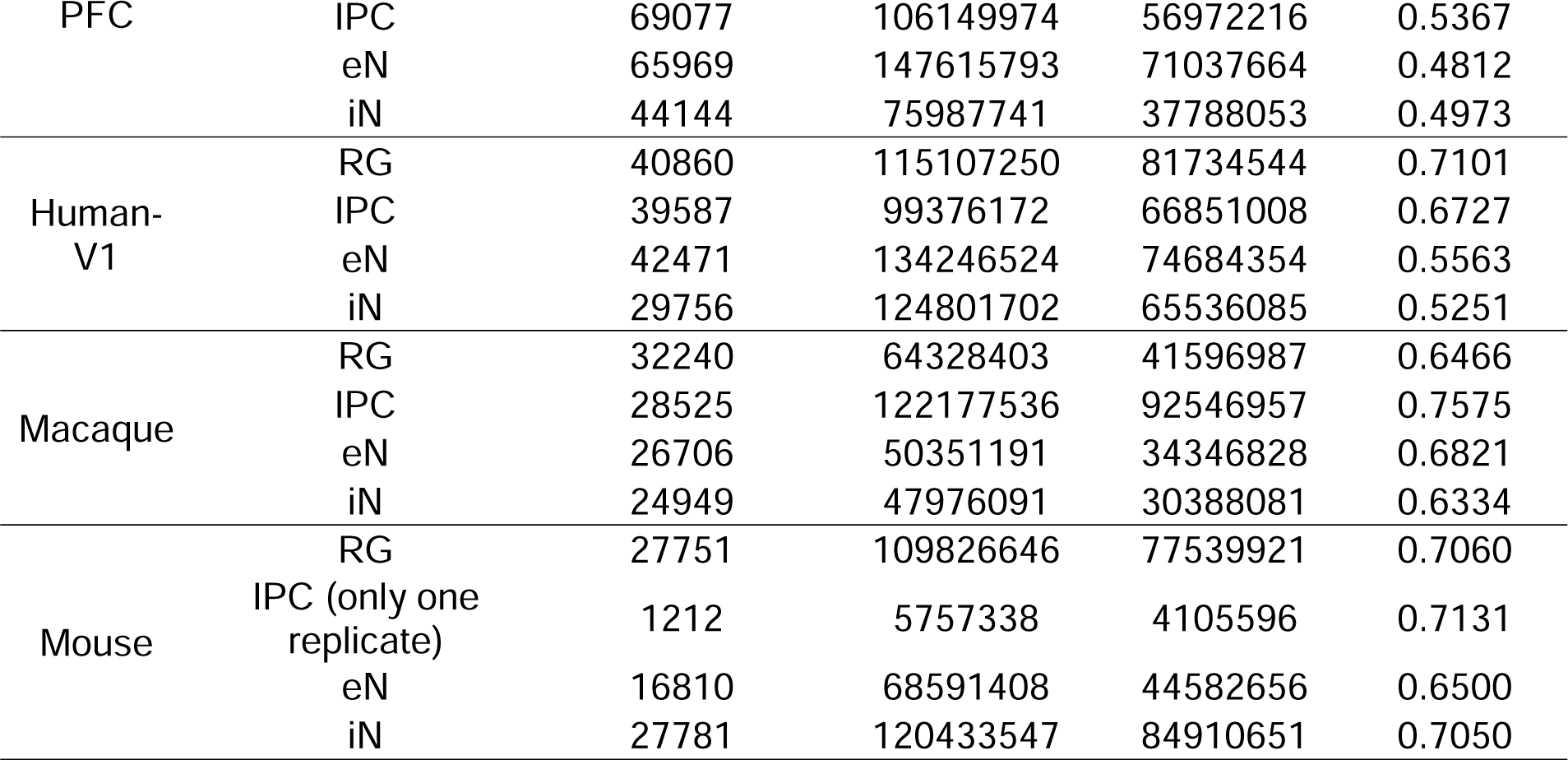
Data samples and library information.

