## Supplemental Figure1-12 for "Long-Range Enhancer Networks Gradually Emerge as Key Regulators During Human Cortical Neurogenesis"

Figure S1. Gel analysis of determining the optimal concentration of DNase I in SCOPE-C library construction and gel cutting strategy for sequencing.

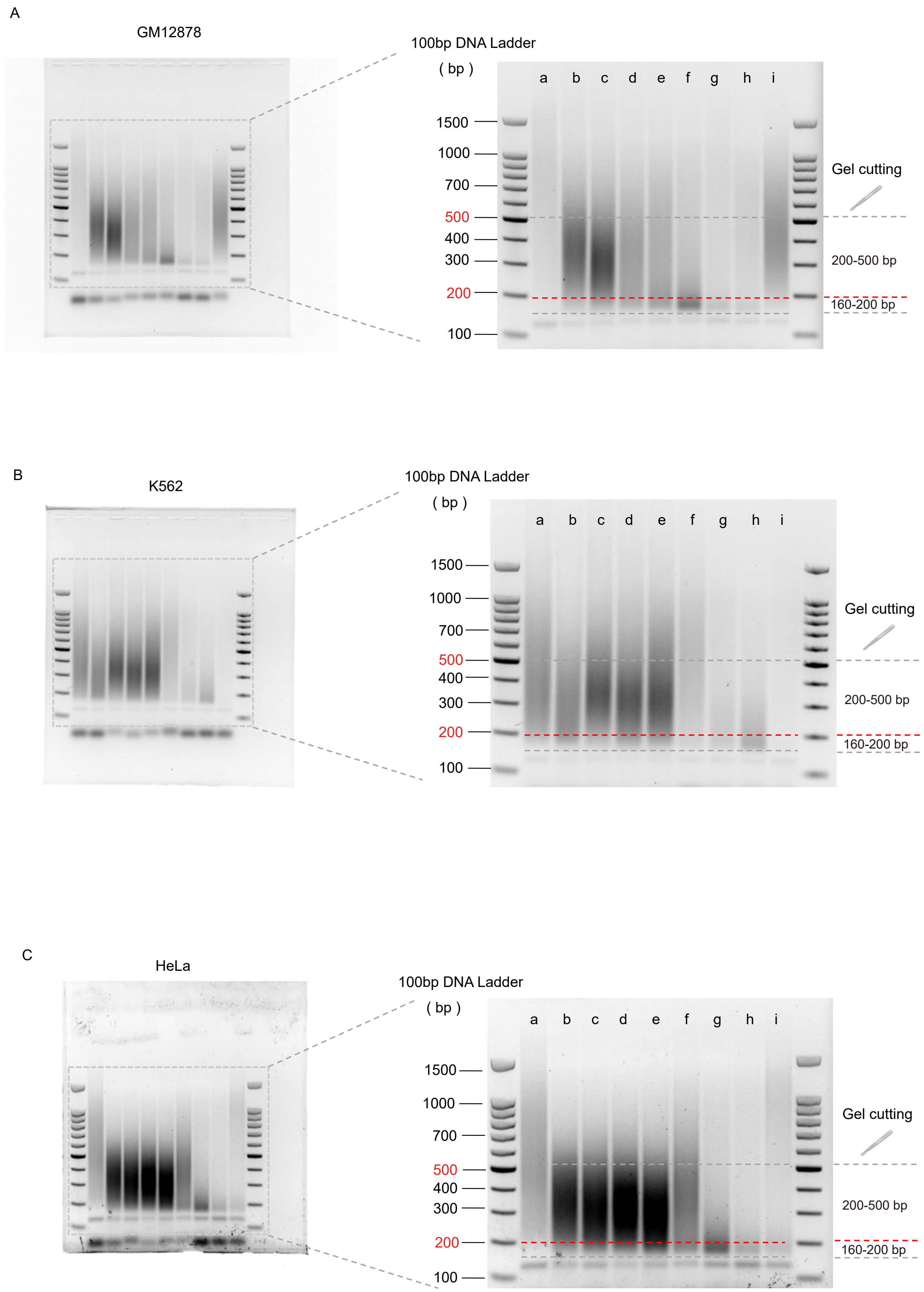

Figure S2. Determining the optimal concentration of DNase I in SCOPE-C experiments.

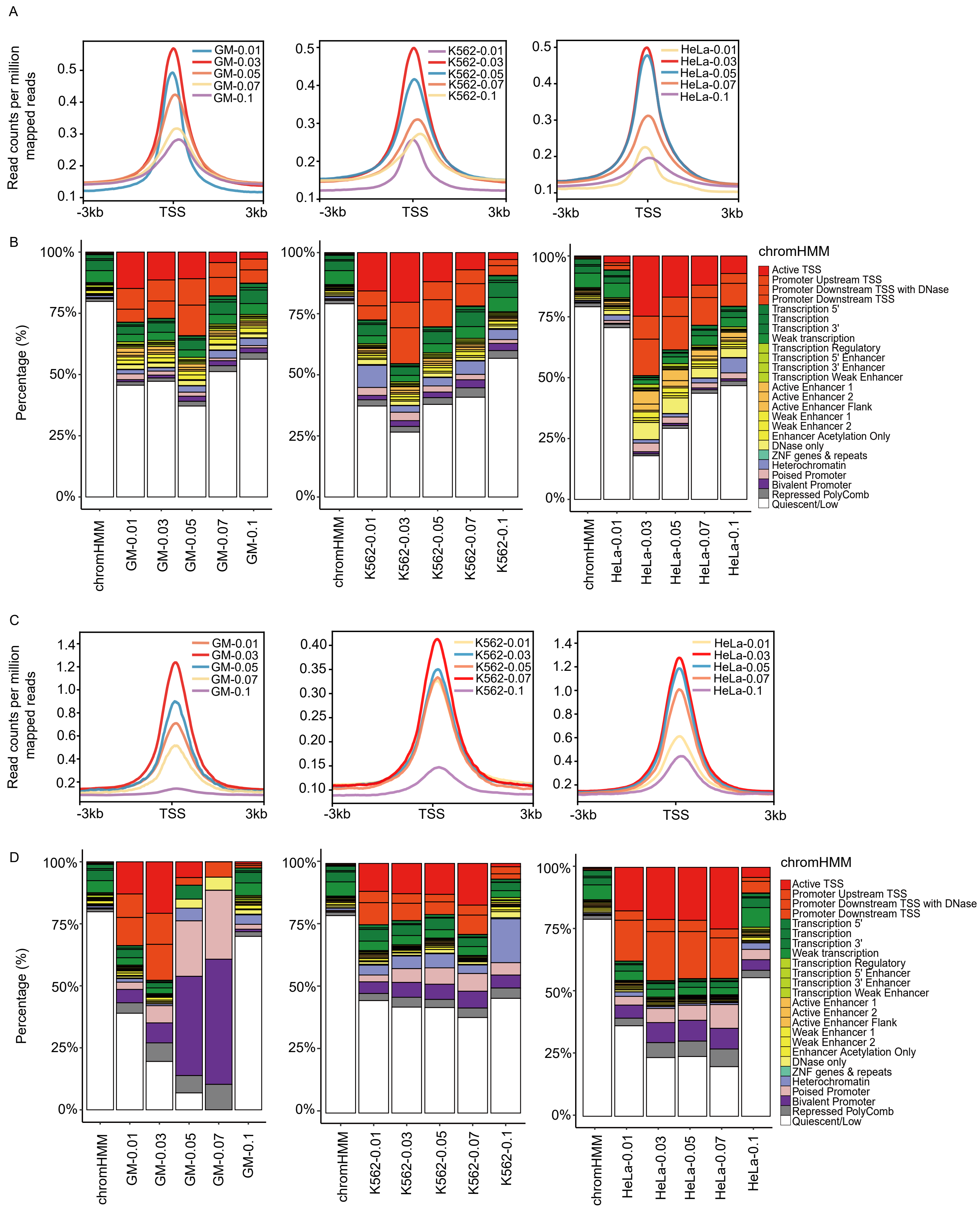

Figure S3. SCOPE-C accurately identifies DHS open-chromatin peaks in bulk/mini-bulk/single cells.

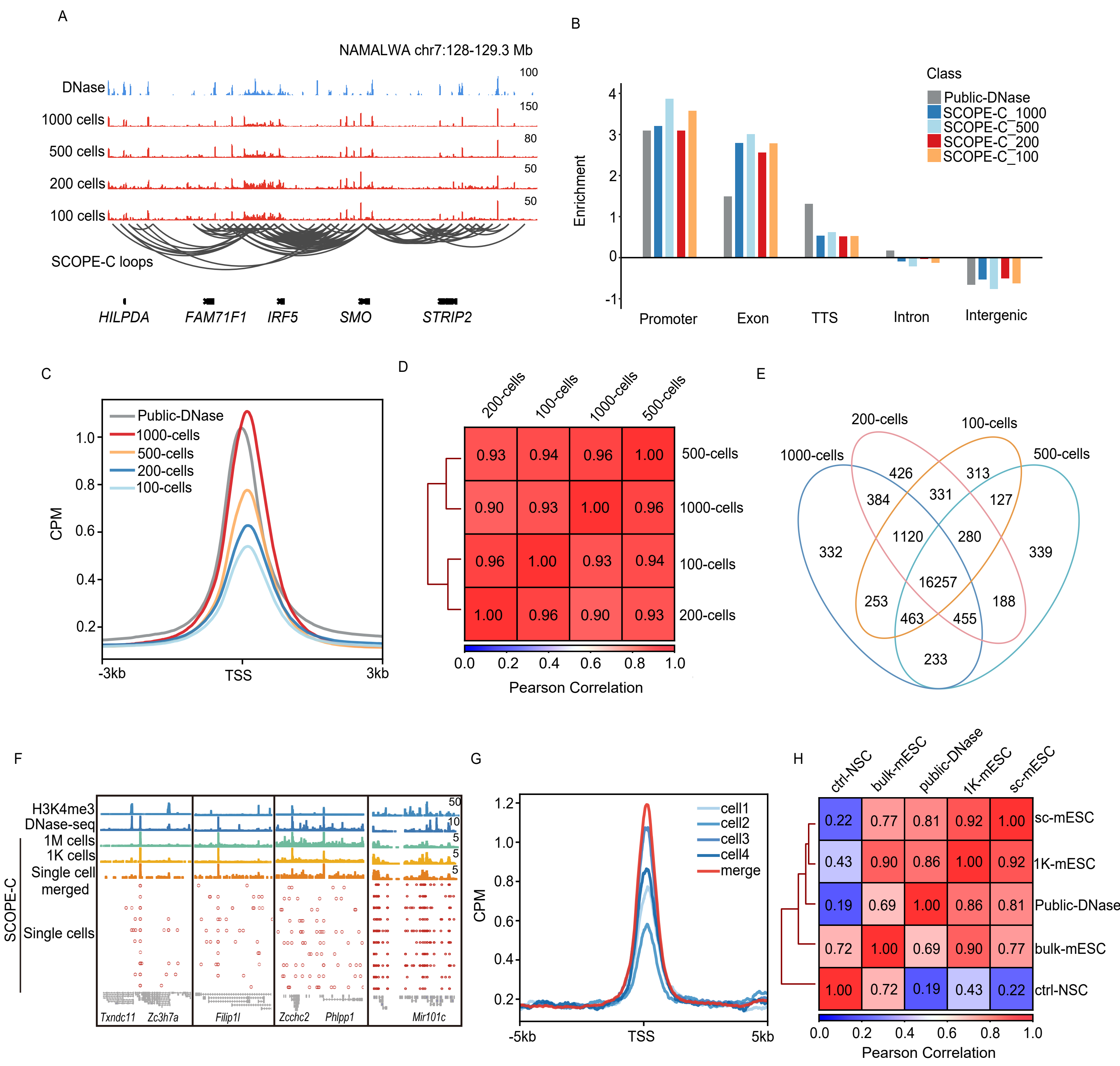

Figure S4. SCOPE-C faithfully capture the key feature of genome organization in bulk/mini-bulk/single cells.

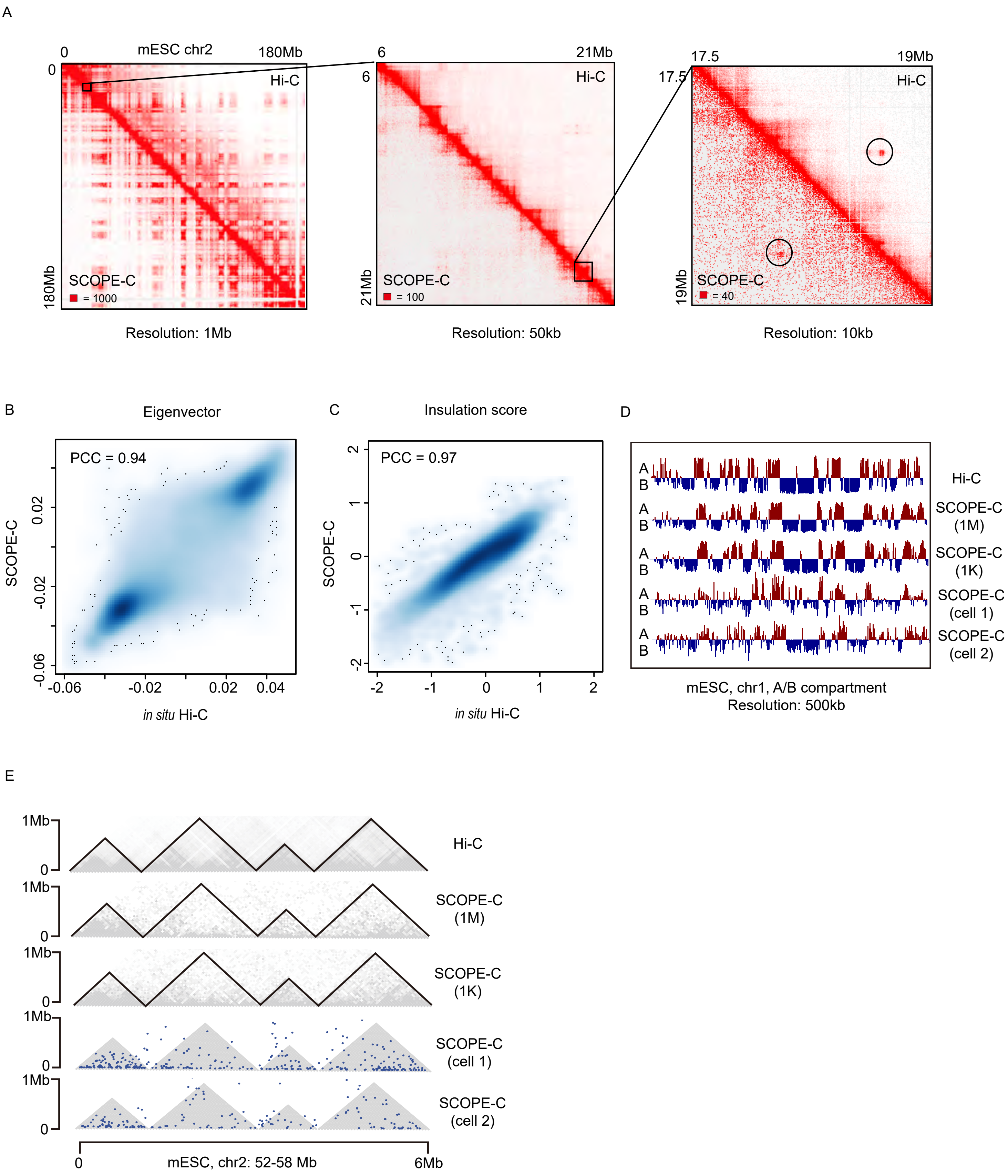

Figure S5. Representative contour plots depicting FACS gating strategy.

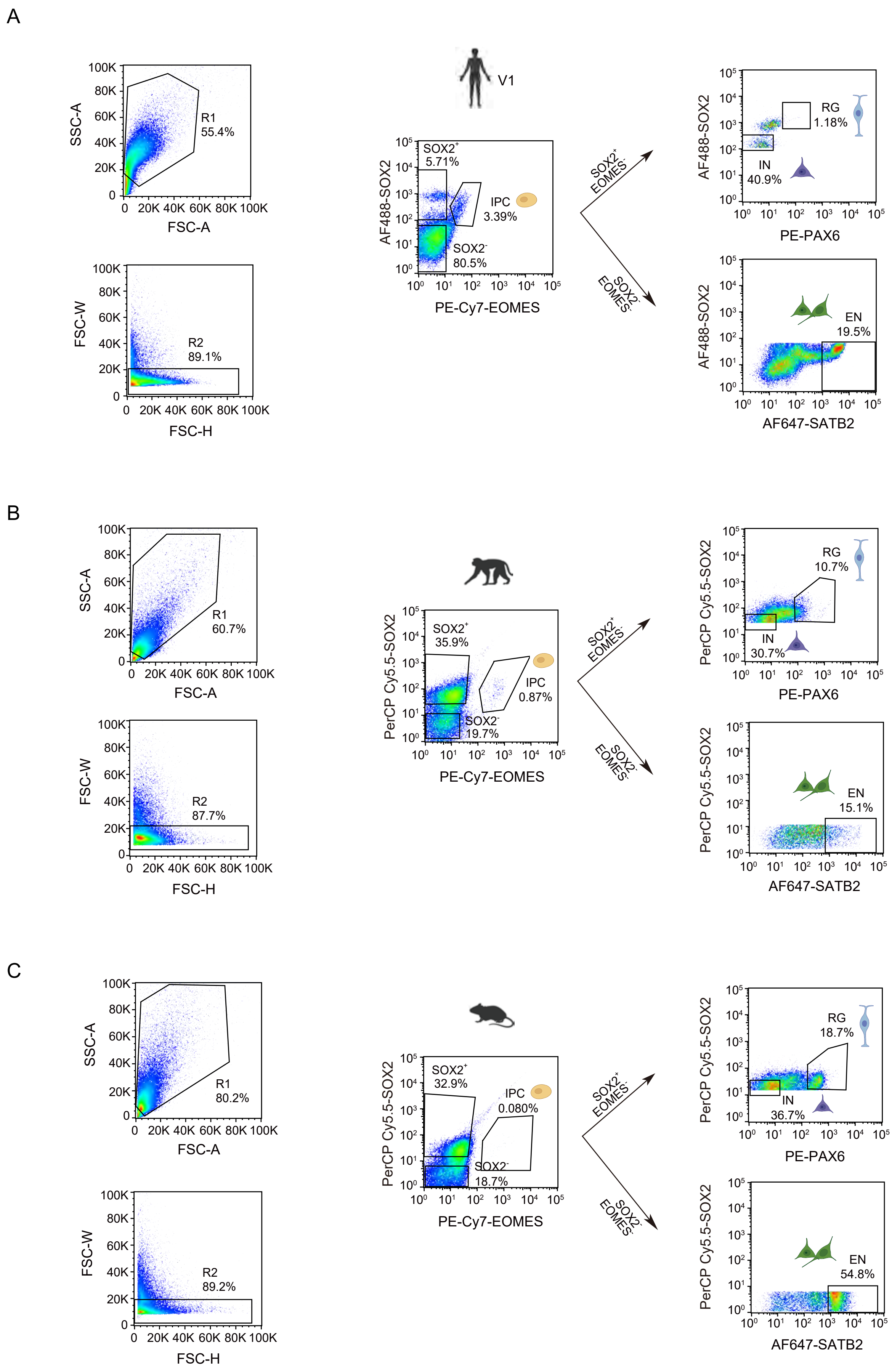

Figure S6. Quality control results for human fetal brain (PFC) SCOPE-C data.

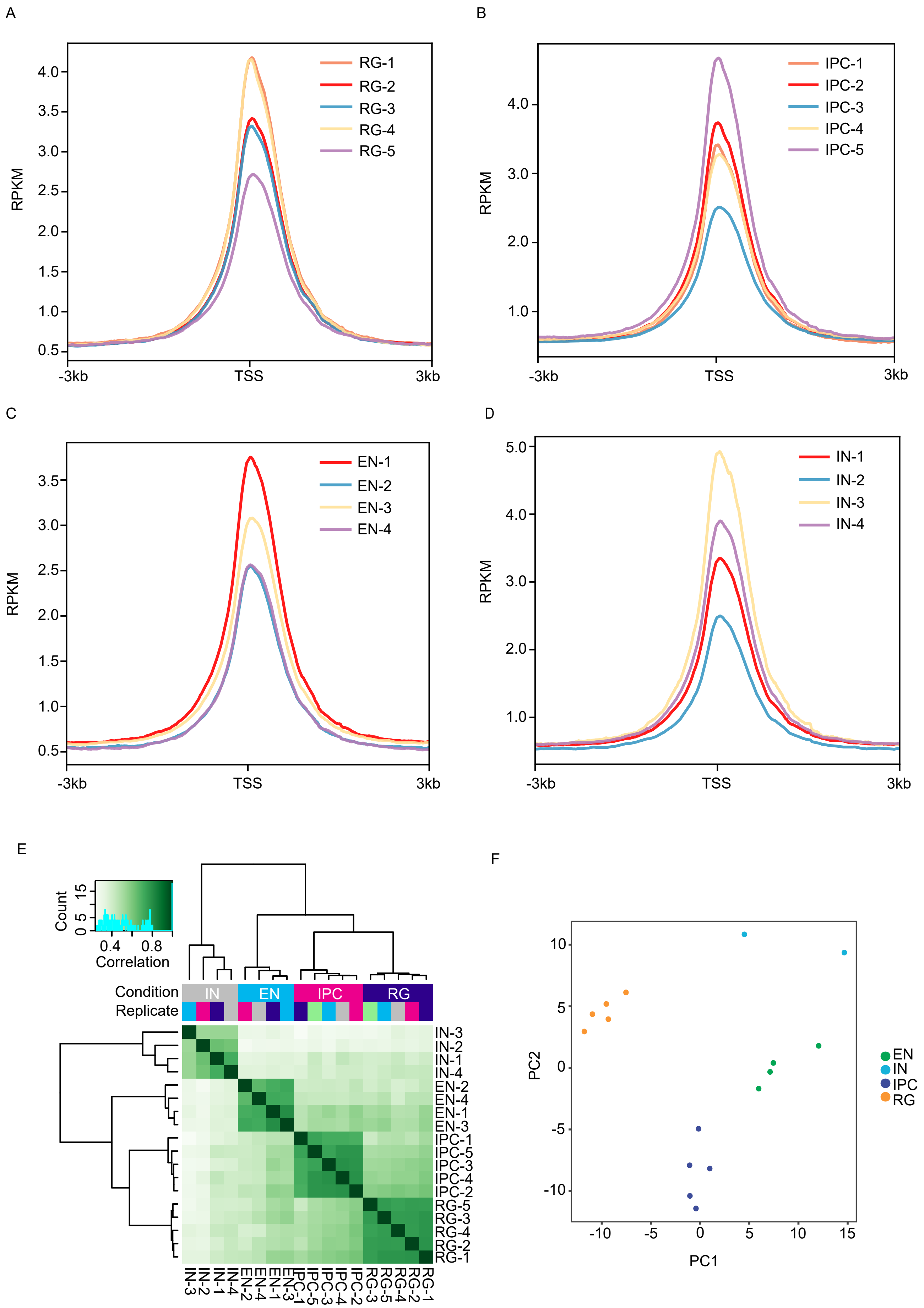

Figure S7. Quality control results for human fetal brain (V1) SCOPE-C data.

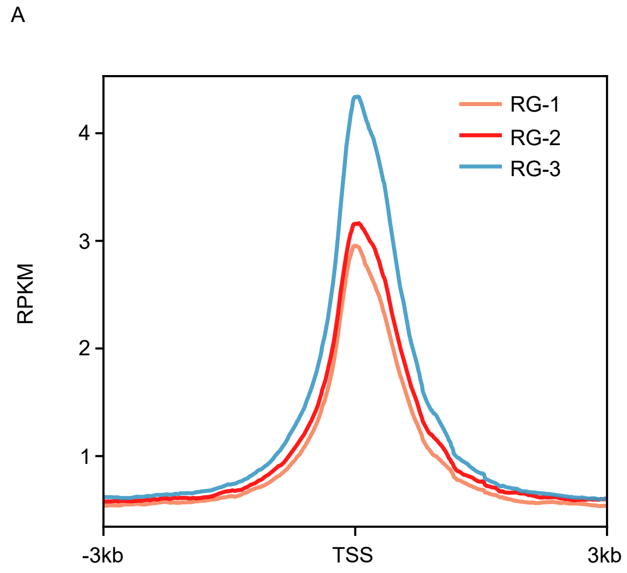


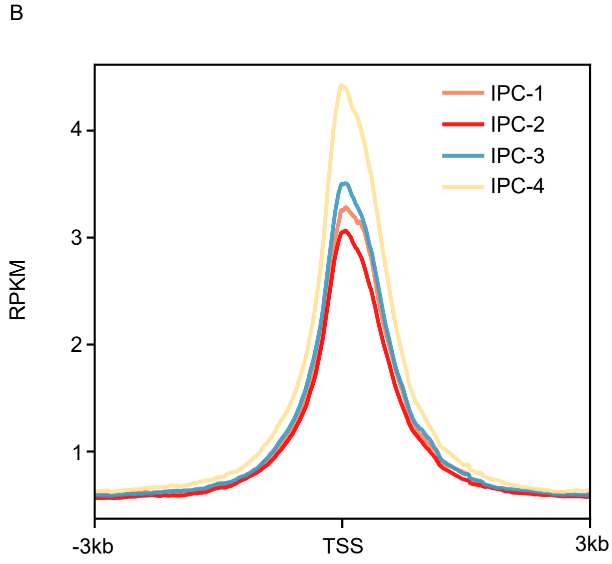


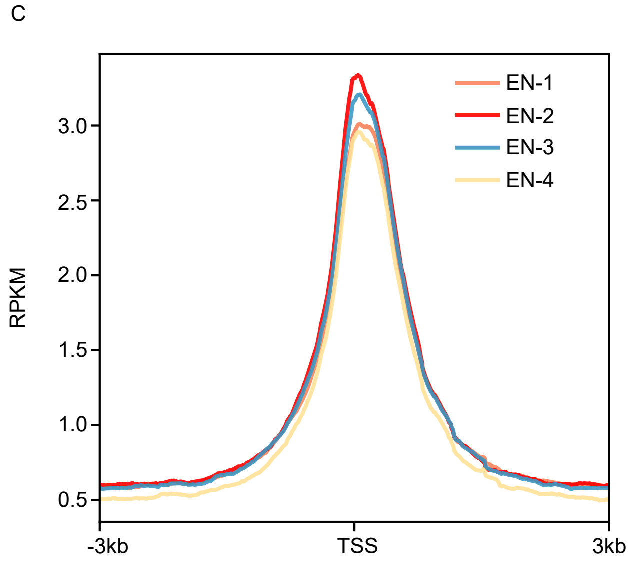


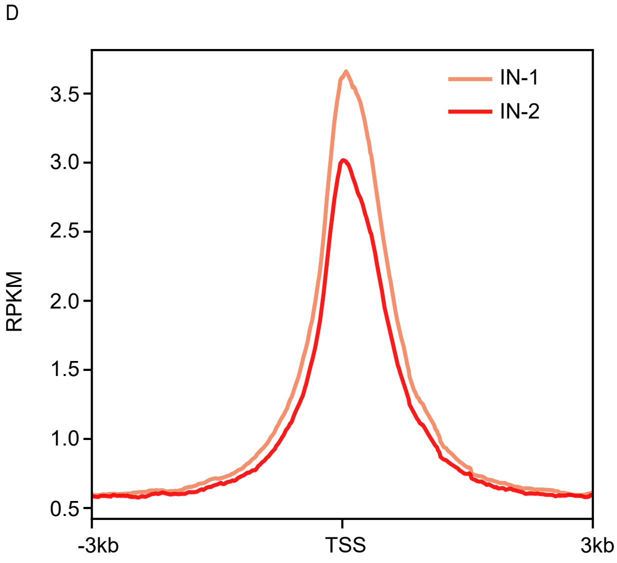


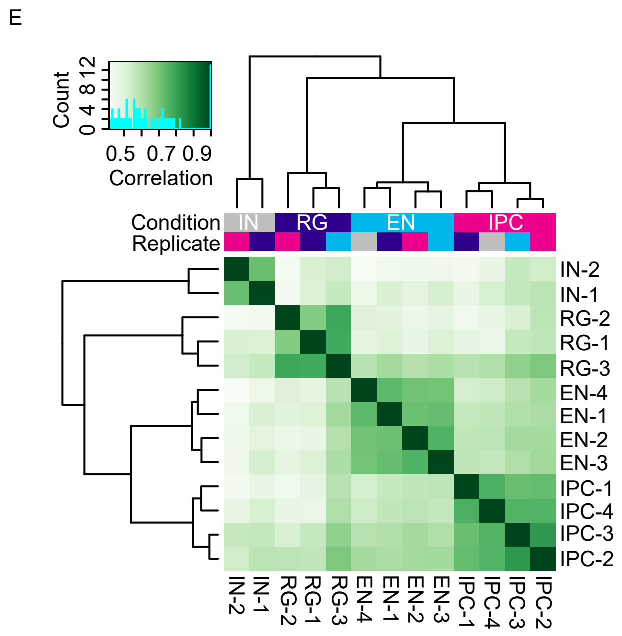


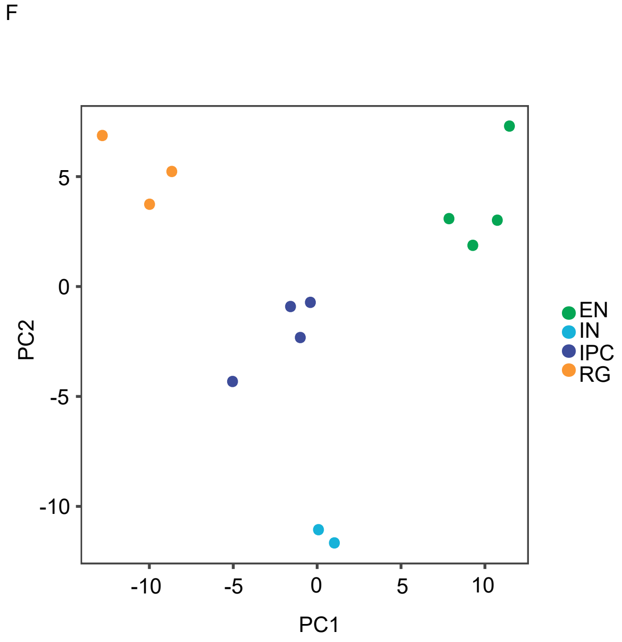

Figure S8. Comparison of human prefrontal cortex (PFC) and primary visual (V1) open chromatin signals.

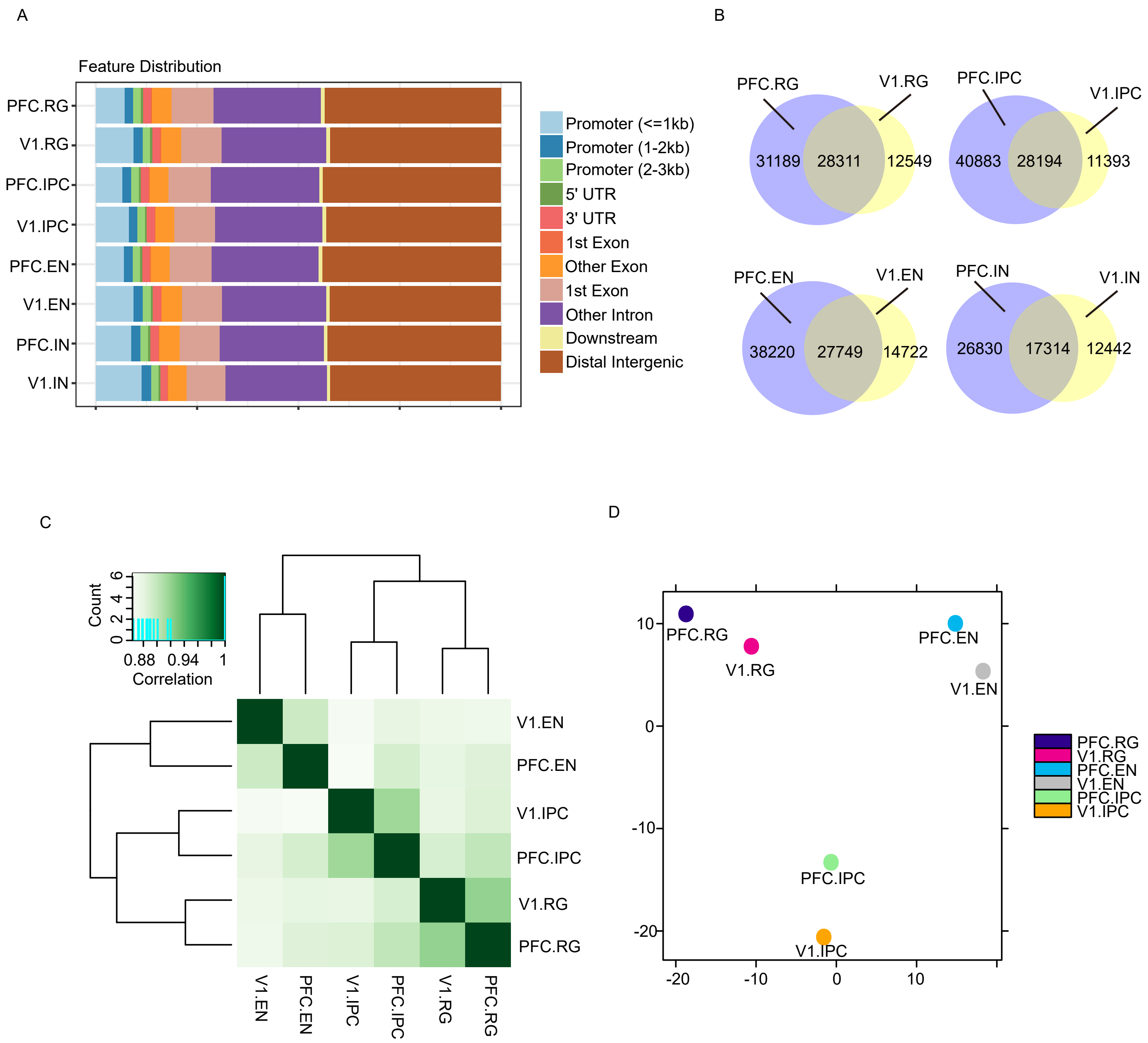

Figure S9. The chromatin openness at HOCl-regulated gene promoters in SCOPE-C and ATAC-seq data.

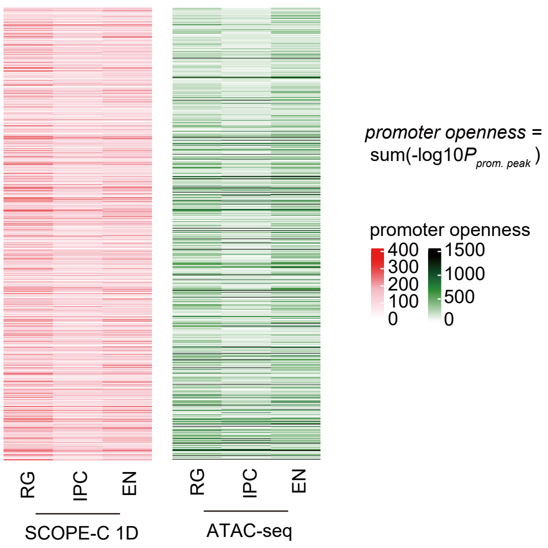

Figure S10. Quality control results for macaque fetal brain SCOPE-C data.

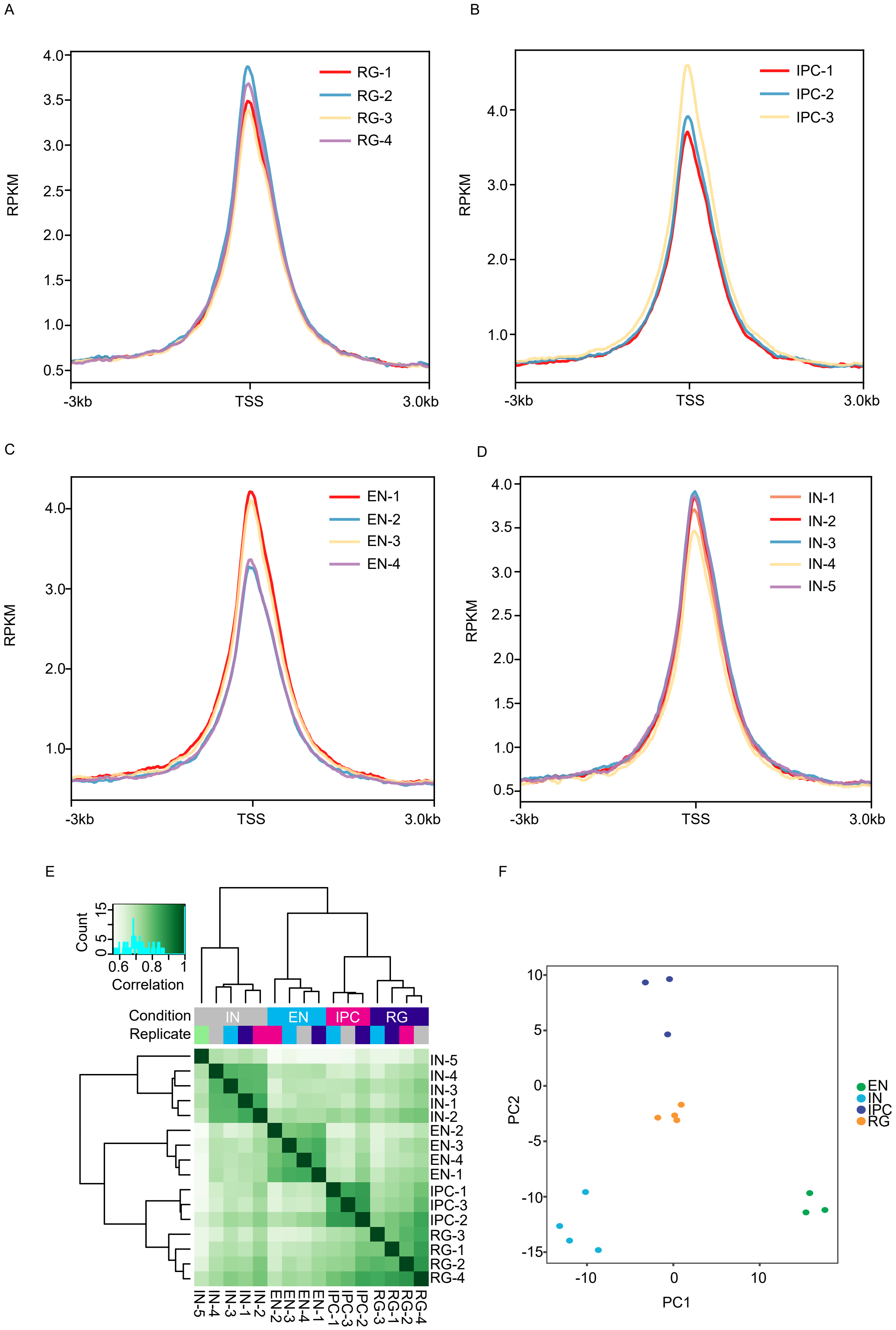

Figure S11. Quality control results for mouse fetal brain SCOPE-C data.

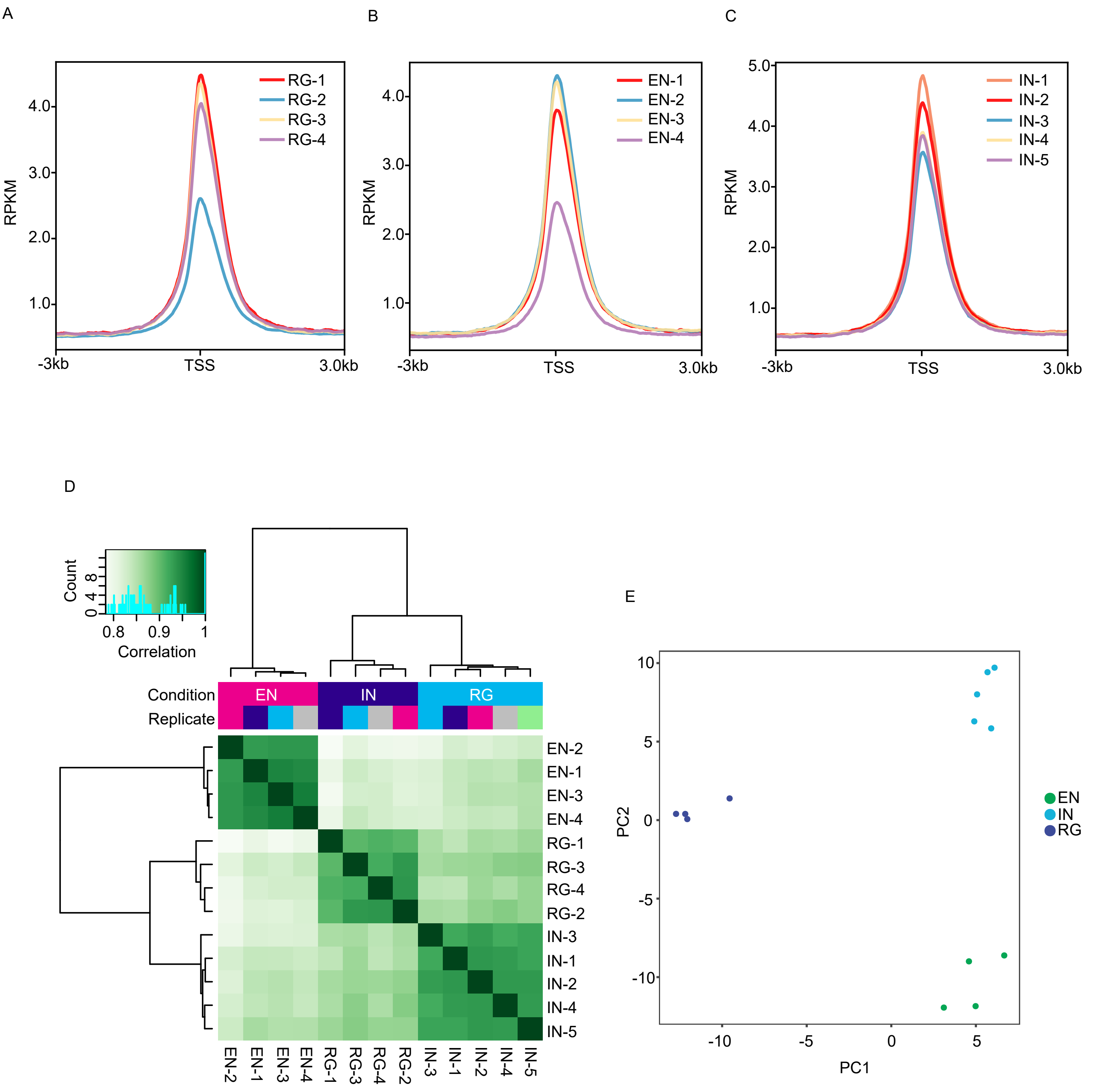

Figure S12. Principal component analysis results of "meta-cell" datasets composed of two groups of single-cell transcriptome data derived from human and mouse.

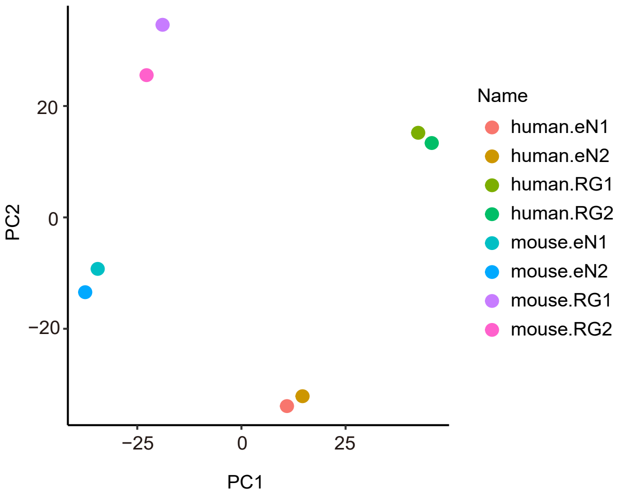
